# “Tumor-to-endothelium mitochondrial transfer licenses endothelial cells for CD8^+^ T cell recognition via mitochondrial neoantigen presentation”

**DOI:** 10.64898/2026.01.30.702567

**Authors:** Francesca Costabile, Pierini Stefano, Renzo Perales-Linares, Nektaria Leli, Adham Bear, Cameron J. Koch, Beatriz M. Carreno, Michael Lotze, Lerry Singh, Marny Falck, Andrea Facciabene

## Abstract

Renal cell carcinoma (RCC) frequently exhibits resistance to immune checkpoint blockade, highlighting the need for strategies that enhance tumor-specific T cell priming and improve immune access to the tumor microenvironment. Here we show that vaccination targeting tumor-associated mitochondrial antigens (TAMAs), derived from tumor-specific mitochondrial DNA (mtDNA) missense mutations, synergizes with PD-1/PD-L1 blockade to overcome checkpoint refractoriness in the RENCA RCC model. TAMAs vaccination elicits antigen-specific T cell responses, increases intratumoral CD8^+^ T cell infiltration, and reduces immunosuppressive myeloid populations, resulting in delayed tumor progression and improved survival when combined with checkpoint inhibition. In parallel, TAMAs + checkpoint blockade induces vascular remodeling characterized by increased pericyte coverage, reduced vascular leakage, improved perfusion and reduced hypoxia. Mechanistically, vascular remodeling is driven by CD8^+^ T cell–dependent, IFN_γ_-associated immune activity and is associated with endothelial apoptosis and diminished intratumoral CD31 signal. We further identify tumor-to-endothelium mitochondrial transfer as a mechanism linking mitochondrial neoantigens to the tumor vascular compartment: tumor-derived mitochondria enter human and mouse endothelial cells *in vitro* and *in vivo*, and tumor-associated mtDNA mutations are detectable in endothelial fractions from murine tumors and human RCC specimens. Human endothelial cells can present mitochondrial neoantigens via MHC class I and become targets of TAMAs-specific CD8^+^ T cell cytotoxicity, including following mitochondrial acquisition from tumor cells. Together, these findings establish mitochondrial neoantigen immunity as a tractable approach to enhance checkpoint responses and reveal mitochondrial transfer as an antigenic bridge that expands immune targeting to the tumor vasculature.

## Results

Tumor-associated antigens represent the *“sine qua non”* requirement for durable T cell– mediated cancer control. Innate immune pathways, including dendritic cells activation, type I interferon signaling and NK cells cytotoxicity, are essential for antigen sensing, amplification, and effector execution; however, sustained tumor regression ultimately depends on antigen-specific adaptive immunity. Yet truly cancer-specific and highly immunogenic antigens remain relatively rare, and the number of high-quality neoantigens available for effective T cell priming is often limited. This remains a major bottleneck for generating robust and long-lasting antitumor T cell responses. Accordingly, identifying new and reliable sources of immunogenic tumor antigens remains a central goal for next-generation cancer immunotherapy^1,2^.

Mitochondrial DNA (mtDNA) mutations may provide such a source^3–6^. Tumors can accumulate mtDNA missense mutations due to elevated oxidative stress, the close proximity of mtDNA to the respiratory chain, and relatively limited repair capacity compared with nuclear DNA. These mutations can alter amino acids within oxidative phosphorylation (OXPHOS) proteins, potentially impacting respiration, while also generating mitochondrial neoantigens. In parallel, mitochondria retain features of their bacterial ancestry, such as the generation of N-formylated peptides and CpG motifs, that can function as innate danger signals. Upon mitochondrial stress or release of mitochondrial contents, formylated peptides can engage formyl peptide receptors on myeloid cells, promoting inflammatory activation, dendritic cells maturation, and antigen presentation. Thus, mitochondrial-derived signals may serve as endogenous adjuvants that enhance the immunogenicity of mitochondrially encoded tumor antigens, while durable antitumor efficacy remains critically dependent on antigen-specific adaptive immunity^7–12^.

Consistent with this concept, recent clinical and mechanistic studies have shown that tumor mtDNA mutational burden can associate with improved responses to immune checkpoint blockade, supporting a functional link between mitochondrial genetic alterations, tumor state, and sensitivity to T cell–mediated immunotherapy^13^.

In earlier work, we identified recurrent missense mtDNA mutations in the RENCA murine renal carcinoma model and generated a dendritic cells (DCs) vaccine incorporating Tumor Associated Mitochondria Antigens (TAMAs). Although this strategy rejected the tumor in prophylactic setting while slowing down tumor progression in therapeutic settings, tumors ultimately escaped immune control^3,14^. Given that immune checkpoint inhibitors (ICIs) are a mainstay of treatment in renal cell carcinoma (RCC), and that PD-1/PD-L1 blockade can reinvigorate exhausted T cells^15,16^, we hypothesized that combining TAMAs vaccination with checkpoint blockade could amplify and prolong TAMA-specific CD8^+^ T cell immunity, thereby improving tumor control.

Here we show that TAMAs vaccination synergizes with PD-1/PD-L1 blockade in RENCA, resulting in enhanced tumor control, increased CD8^+^ T cell infiltration, and reduced accumulation of immunosuppressive myeloid populations within the tumor microenvironment. Notably, this combinatorial strategy also induced marked vascular remodeling, including increased pericyte coverage, reduced vascular leakage, improved perfusion, and decreased hypoxia, features consistent with vascular normalization^17^. Because effective CD8^+^ T cells infiltration depends on functional vascular support, these observations suggested that TAMAs-driven immunity may impact not only malignant cells but also stromal compartments that regulate immune entry and function^18^.

At the same time, intercellular mitochondrial transfer has emerged as a widely observed form of cellular communication in both physiological and pathological contexts. In cancer, most studies have focused on mitochondria being transferred into tumor cells, where they can enhance metabolic fitness, facilitate adaptation to stress, or promote resistance to therapy. By contrast, far less is known about mitochondria moving in the opposite direction, out of tumor cells and into stromal populations and essentially almost nothing is understood regarding how such transfer might shape antitumor immunity^19–21^.

Our data now demonstrate that tumor cells transfer mitochondria to endothelial cells *in vivo*, and that the transferred mitochondria carry the same mtDNA missense mutations that define TAMAs. Endothelial cells acquiring tumor mitochondria display tumor-derived mtDNA mutations at detectable heteroplasmic levels and can be recognized and killed by TAMAs-specific CD8^+^ T cells, particularly in the context of checkpoint blockade. These findings identify mitochondrial transfer as an unexpected “antigen bridge” between malignant cells and the tumor vasculature, effectively converting tumor endothelial cells into additional immunological targets.

Finally, we establish the clinical relevance of this mechanism in human RCC. Across patient cohorts, we identified tumor-specific mtDNA missense mutations and detected the same mutations within endothelial-enriched tumor compartments, consistent with tumor-to-endothelium mitochondrial transfer in situ. We further show that multiple human TAMAs peptides elicit strong IFN_γ_ responses from primed CD8^+^ T cells, and that endothelial cells can present mitochondrial neoantigens via MHC-I in a manner sufficient to trigger cytotoxic recognition. In parallel, analyses integrating mtDNA mutation profiling with quantitative immunostaining revealed vascular phenotypes associated with recurrent mitochondrial gene mutations, and multiplex imaging demonstrated apoptotic CD31^+^ vascular structures in close spatial association with CD8^+^ T cells in human tumors, findings consistent with immune-linked endothelial injury.

Together, these data establish mitochondrial neoantigens as an underappreciated class of targets for cancer immunotherapy and uncover a mechanism by which tumor-to-endothelium mitochondrial transfer broadens the cellular target landscape of cytotoxic CD8^+^ T cells to include the tumor vasculature. By enabling stromal antigen presentation and immune-mediated vascular remodeling, this process may contribute to overcoming resistance to checkpoint blockade in solid tumors. Because mtDNA mutations are patient- and tumor-specific, TAMAs may also provide a rational substrate for personalized neoantigen vaccination strategies.

### 1. TAMAs vaccination synergizes with PD-1/PD-L1 blockade to enhance antitumor immunity in RCC

We first assessed whether our mitochondrial antigen TAMAs–based vaccination strategy could cooperate with ICI in the RENCA renal cell carcinoma model. RENCA tumors displayed significantly increased PD-L1 expression relative to normal kidney, as demonstrated by immunofluorescence staining and quantification of PD-L1 signal intensity (Fig. 1A).

**Figure 1.**
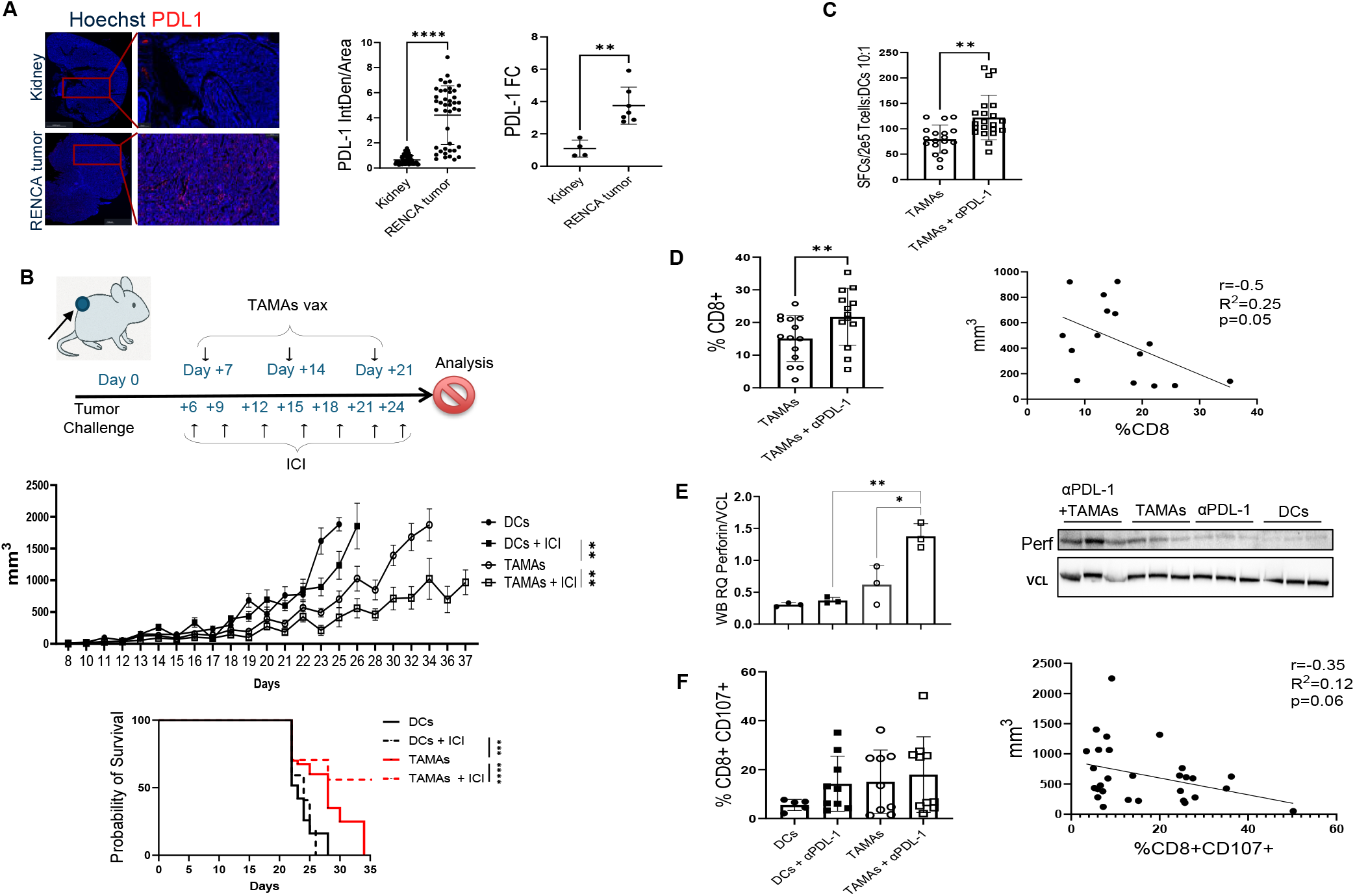
TAMAs vaccination enhances response to checkpoint blockade in RENCA RCC. (A) Representative immunofluorescence images showing PD-L1 expression in naïve kidney and RENCA tumors (Hoechst, nuclei; PD-L1, red). IF and qPCR quantification of PD-L1 signal is shown on the right. (B) From top to bottom, schematic of RENCA tumor implantation and treatment schedule, longitudinal tumor growth monitoring and endpoint analyses. (C) IFN_γ_ ELISPOT responses of CD8^+^ T cells to BMDCs pulsed with TAMAs peptides and/or RENCA mitochondrial lysate. Peptide- and lysate-driven responses are shown as the summed response (spot-forming cells, SFC, per indicated number of CD8^+^ T cells). (D) Flow cytometric quantification of intratumoral CD8^+^ T cells across treatment groups. Right, correlation between tumor volume and CD8^+^ T cell abundance. (E) Immunoblot showing perforin expression in tumors across treatment groups (VCL loading control) with densitometric quantification. (F) Flow cytometric analysis of activated CD8^+^ T cells (CD107a^+^) across treatment groups and correlation with tumor volume. Data are shown as mean ± s.e.m. unless otherwise indicated. Statistical comparisons were performed using one-way ANOVA with multiple-comparison correction.

BALB/c mice were injected subcutaneously with 1×10□RENCA cells and, six days later, randomized to receive one of four treatment regimens: (a) control DC vaccine ± ICI, or (b) TAMAs vaccine ± ICI. TAMAs were delivered subcutaneously on days 7, 14, and 21, whereas anti–PD-L1/anti–PD-1 antibodies were administered intraperitoneally following the schedule shown in Fig. 1B.

TAMAs vaccination alone significantly delayed tumor progression and reduced tumor volume compared with control mice (Fig. 1B). Notably, the TAMAs + ICI combination further improved tumor control relative to TAMAs alone, resulting in significantly smaller tumors and extended survival (Fig. 1B). In contrast, ICI alone did not improve tumor growth kinetics compared with DC controls (Fig. 1B), indicating limited activity of checkpoint blockade in this model in the absence of vaccination^22,23^.

We next assessed immune activation in response to TAMAs vaccination. TAMAs induced T-cell reactivity against BMDCs pulsed either with RENCA mitochondrial lysate or peptides derived from the recurrent COX1 mtDNA mutation (Fig. 1C). In line with increase of vaccination potency, at the tumor site, TAMAs + ICI increased the frequency of intratumoral CD8^+^ T cells compared with TAMAs alone (Fig. 1D; Supplementary Fig. 1A). Across individual tumors, the correlation between CD8^+^ T-cell frequency and tumor size showed a negative correlation (Fig. 1D). In parallel, perforin protein abundance was increased in tumors treated with TAMAs + ICI compared with other groups (Fig. 1E). The frequency of CD8^+^CD107a^+^ T cells also showed a trend toward a negative correlation with tumor volume (Fig. 1F).

Consistent with enhanced antitumor activity, intratumoral CD8^+^ T-cell abundance positively correlated with TUNEL^+^ apoptotic nuclei within the tumor (Supplementary Fig. 1B). In contrast, no significant correlations were observed between CD4^+^ T-cell infiltration and tumor size or tumor apoptosis (Supplementary Fig. 1C). Combination of TAMAs + ICI also significantly reduced the frequency of CD11b^+^Gr1^+^ myeloid cells relative to control conditions (Supplementary Fig. 1D).

Together, these data demonstrate that TAMAs vaccination enhances antitumor immunity in RENCA tumors and that combination with checkpoint blockade further improves tumor control, in association with increased intratumoral CD8^+^ T cell frequency and cytotoxic effector signatures.

### 2. TAMAs vaccination and ICI cooperate to promote vascular remodeling consistent with vessel normalization

Given the requirement of functional vasculature for T cells infiltration and the increased intratumorally CD8^+^ T-cell infiltration we observed with TAMAs + ICI, we next investigated whether combination treatment modulated the tumor vasculature^24,25^. Doppler ultrasound analysis showed that tumors treated with TAMAs + ICI exhibited a significantly higher perfusion index compared with controls (Fig. 2A; Supplementary Fig. 2A). Perfusion index did not correlate with tumor size (Supplementary Fig. 2B), suggesting that perfusion differences were not solely explained by reduced tumor burden.

**Figure 2.**
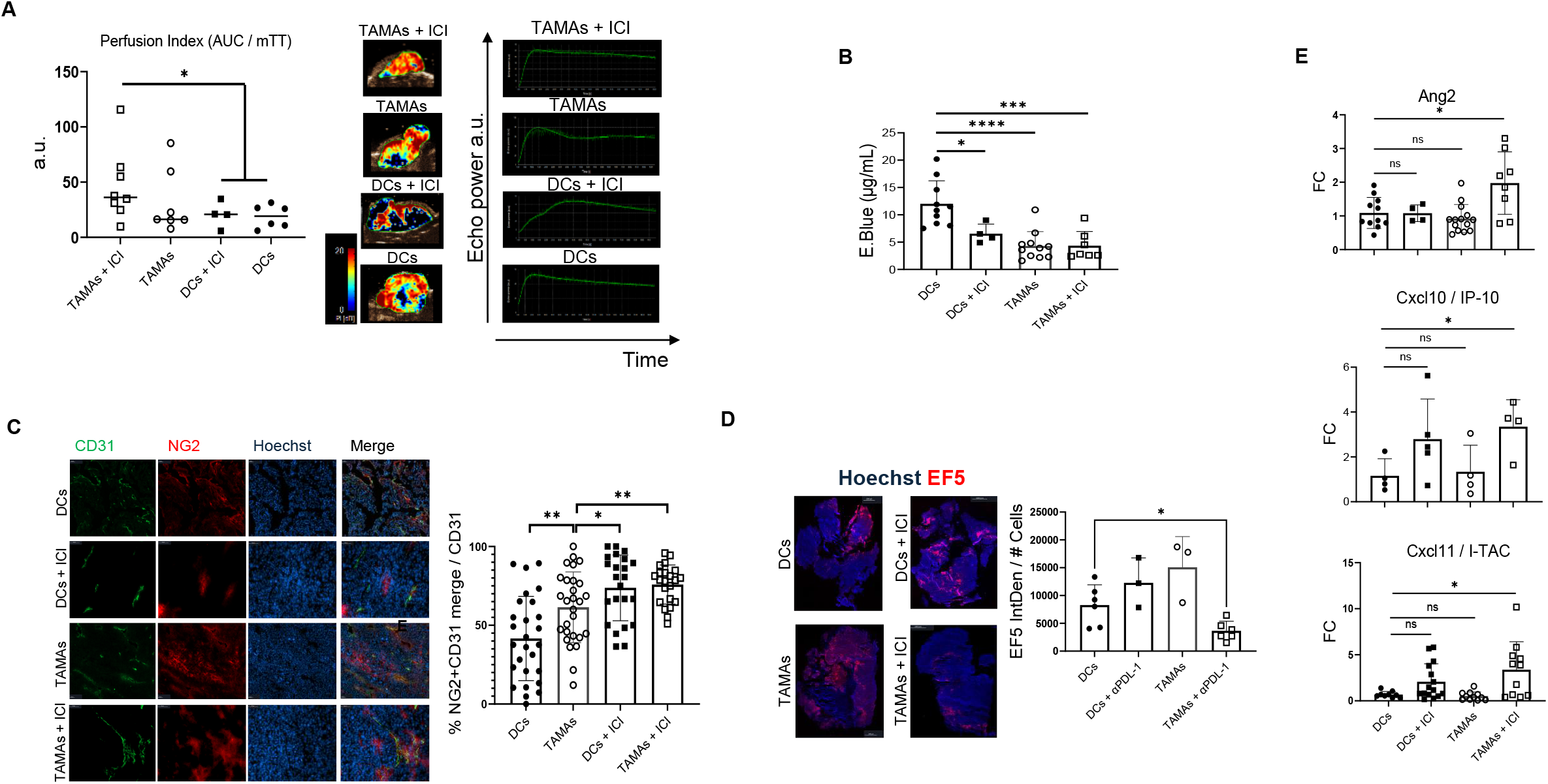
TAMAs + ICI promotes vascular remodeling and improves vessel function in RENCA tumors. (A) Contrast-enhanced ultrasound analysis of tumor perfusion. Left, perfusion index quantification. Middle, representative parametric perfusion maps. Right, representative time– intensity curves. (B) Quantification of Evans Blue extravasation (vascular leakage) across treatment groups. (C) Representative immunofluorescence images of tumor vasculature (CD31) and pericyte coverage (NG2) with quantification of NG2^+^ coverage normalized to CD31^+^ area. (D) Representative EF5 staining for hypoxia and quantification of EF5^+^ signal. (E) qPCR analysis of vascular/immune-related transcripts in tumors (Ang2, Cxcl10/IP-10, Cxcl11/I-TAC; fold change vs control). Each dot represents one mouse/tumor. Data are mean ± s.e.m. Statistical significance was assessed using one-way ANOVA with multiple-comparison correction.

Evans Blue^26^ quantification demonstrated decreased vascular leakage in treated groups, with the most pronounced reduction observed in the TAMAs + ICI group (Fig. 2B). Immunofluorescence staining for NG2 (pericytes) and CD31 (endothelium) revealed an increased pericyte coverage across treatment arms^27^, including a significant increase in the combination group (Fig. 2C). EF5 staining, in line with better tumor blood perfusion, further indicated reduced intratumoral hypoxia in the TAMAs + ICI group compared with controls (Fig. 2D)^28^.

To probe molecular mediators associated with vascular remodeling, we measured expression of cytokines and chemokines linked to endothelial activation and immune recruitment. TAMAs + ICI increased tumor expression of Ang2, Cxcl10, and Cxcl11 relative to control conditions^29^ (Fig. 2E).

Together, these data indicate that TAMAs vaccination and ICI promote vascular remodeling characterized by increased pericyte coverage, reduced leakage, improved perfusion and reduced hypoxia, features consistent with vascular normalization^17^.

### 3. CD8^+^ T cells and IFN_γ_ are required for vascular remodeling

To define immune programs contributing to tumor vascular remodeling, we performed adoptive T-cell transfer (ACT) of purified CD4^+^ or CD8^+^ T cells in combination with ICI. CD8^+^ ACT + ICI significantly increased αSMA^+^ pericyte coverage on CD31^+^ endothelial structures compared with ICI alone (Fig. 3A). CD4^+^ ACT + ICI also increased pericyte coverage, although with modest magnitude (Fig. 3A). Importantly, IFN_γ_ neutralization reduced pericyte coverage in both transfer settings (Fig. 3A), supporting an IFN_γ_-dependent mechanism of vascular remodeling^27^.

**Figure 3.**
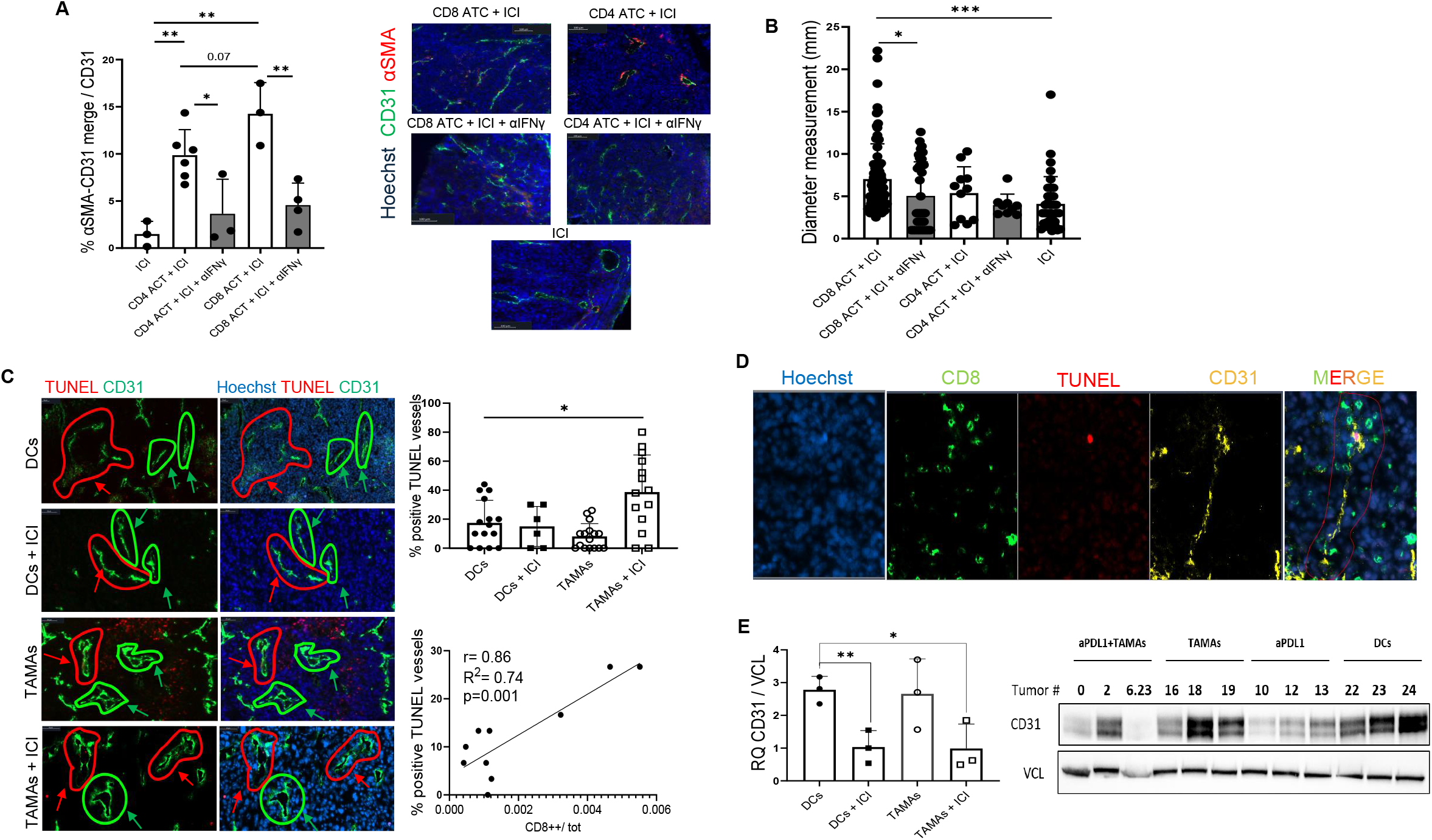
CD8^+^ T cells and IFN_γ_ drive immune-dependent vascular remodeling and endothelial injury. (A) Quantification of αSMA^+^ pericyte coverage normalized to CD31^+^ endothelial area following adoptive transfer (ATC) of CD4^+^ or CD8^+^ T cells in the presence of ICI, with or without IFN_γ_ neutralization. Representative images are shown. (B) Quantification of vessel diameter across indicated treatment/transfer conditions. (C) Representative multiplex immunofluorescence images (CD31, TUNEL) with quantification of apoptotic vessels and correlation with intratumoral CD8^+^ abundance. (D) Representative images showing spatial proximity of CD8^+^ T cells and apoptotic endothelial structures (CD31^+^TUNEL^+^). (E) CD31 protein abundance measured by immunoblot with densitometric quantification. Data are mean ± s.e.m. Statistical comparisons were performed using one-way ANOVA with multiple-comparison correction.

We next quantified vessel diameters as an independent readout of vascular remodeling. CD8^+^ ACT + ICI increased larger vessel diameter count relative to ICI alone, and this effect was abrogated by IFN_γ_ neutralization (Fig. 3B). In contrast, CD4^+^ ACT did not significantly alter vessel diameter (Fig. 3B), suggesting possible distinct effects of CD4^+^ versus CD8^+^ T cells on vascular structure.

We next assessed whether TAMAs vaccination combined with ICI induced endothelial apoptosis. TUNEL/CD31 co-staining demonstrated increased apoptotic endothelial structures in TAMAs + ICI tumors compared with control groups (Fig. 3C). Importantly the frequency of TUNEL^+^ CD31^+^ vessels positively correlated with intratumoral CD8^+^ T-cell abundance (Fig. 3C). *In vivo* Multiplex staining further demonstrated CD8^+^ T cells in proximity to TUNEL^+^CD31^+^ vascular structures (Fig. 3D). Finally, TAMAs + ICI reduced overall CD31 protein abundance relative to control conditions (Fig. 3E), consistent with treatment-associated changes in tumor vascular abundance.

Together, these findings indicate that vascular remodeling in this setting is mostly promoted by CD8^+^ T cells through an IFN_γ_-dependent mechanism and is associated with increased endothelial apoptosis and changes in tumor vascular markers and morphology.

### 4. Tumor-to-endothelium mitochondrial transfer occurs *in vitro, in vivo*, and in human RCC

Intercellular mitochondrial transfer has been increasingly recognized as a component of tumor– stroma communication, but most reports describe mitochondria being imported into tumor cells^30^. Whether mitochondria also move out of tumor cells and into stromal populations *in vivo* remains poorly explored^20^. Because TAMAs arise from tumor-specific mtDNA mutations, we reasoned that any transfer of tumor-derived mitochondria into endothelial cells could deliver these neoantigenic determinants to the vasculature, rendering endothelial cells susceptible to TAMAs-specific CD8^+^ T-cell immunity (Fig. 4A).

**Figure 4.**
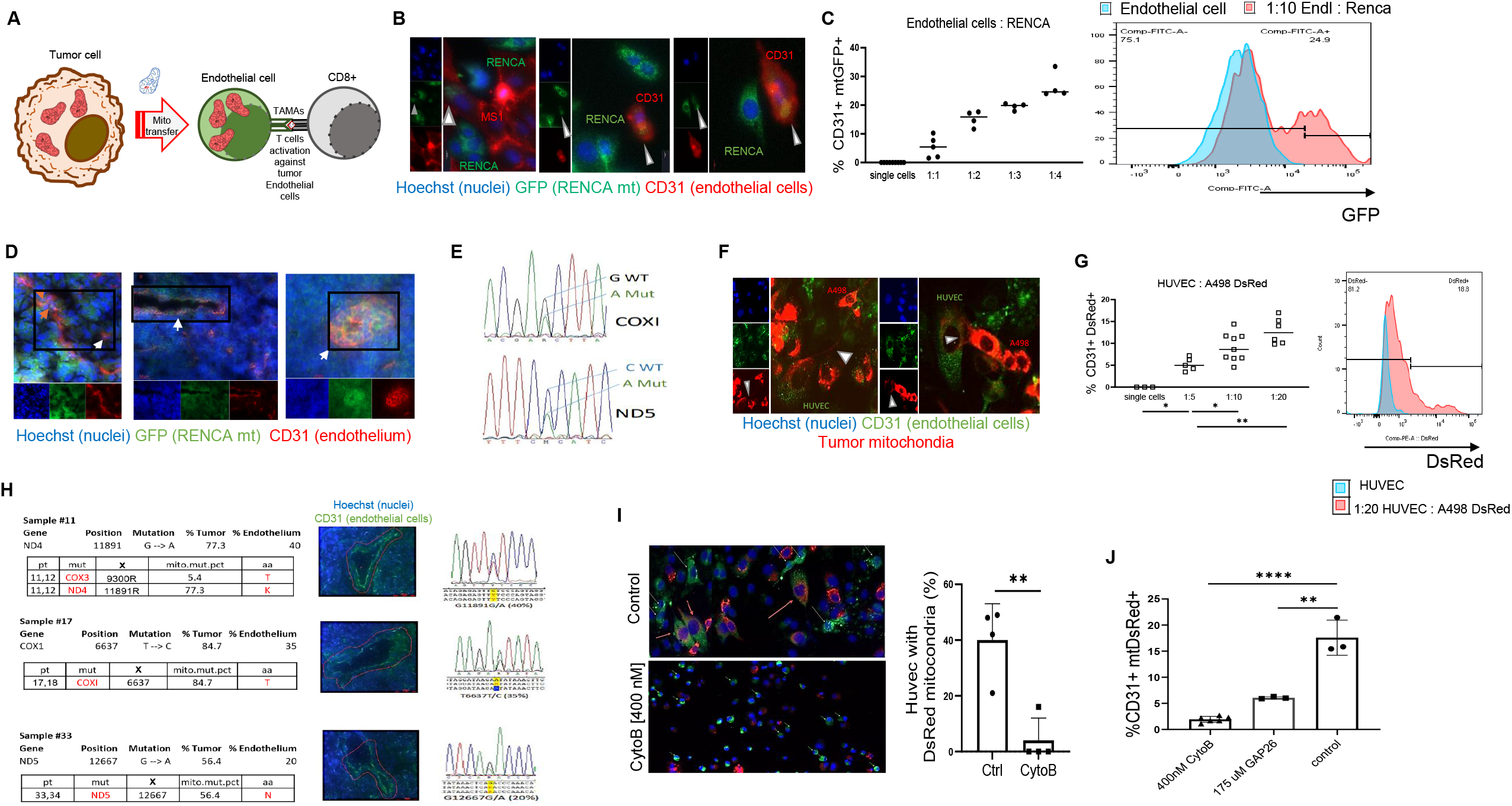
Tumor-derived mitochondria transfer into endothelial cells and tumor-associated mtDNA mutations are detected in endothelial compartments. (A) Working model of tumor-to-endothelium mitochondrial transfer enabling endothelial targeting by mitochondrial neoantigen–specific CD8^+^ T cells. (B) Representative microscopy images showing RENCA mtGFP signal within/adjacent to CD31^+^ endothelial cells *in vitro*. (C) Flow cytometric quantification of transfer to recipient endothelial cells across indicated tumor:endothelial ratios; representative histogram shown. (D) Representative *in vivo* imaging showing mtGFP tumor-mitochondrial signal within CD31^+^ vascular regions. (E) Representative sequencing traces showing tumor-associated mtDNA mutations (COX1, ND5) detected in endothelial purified fractions. (F–G) Human RCC tumor-to-HUVEC mitochondrial transfer visualized by microscopy (F) and quantified by flow cytometry (G). (H) Detection of tumor-associated mtDNA variants in matched RCC tumor and CD31 compartments with representative sequencing traces. (I-J) Pharmacologic inhibition of actin dynamics (cytochalasin B) or connexin signaling (GAP26) reduces mitochondrial transfer to endothelial cells

*In vitro* co-culture of RENCA cells expressing GFP-tagged mitochondria (supplementary Fig 3A) with endothelial cells demonstrated GFP^+^ mitochondrial signal within CD31^+^ endothelial cells by microscopy (Fig. 4B). Flow cytometry confirmed transfer of GFP signal into CD31^+^ endothelial cells and showed that transfer increased as the endothelial:tumor ratio decreased (Fig. 4C).

To test whether this process occurs *in vivo*, we implanted RENCA cells expressing GFP-tagged mitochondria into mice and analyzed established tumors. GFP^+^ mitochondrial structures were detected within CD31^+^ tumor endothelial cells (Fig. 4D). To genetically validate the presence of tumor-associated mtDNA within endothelial compartments, we sequenced mtDNA from highly purified endothelial fractions obtained sequentially by first enrichment endothelial cells purification kit (Miltenyi) followed by cells sorting and detected tumor-associated COX1 and ND5 mutations in the endothelial sample (Fig. 4E).

We next evaluated mitochondrial transfer in a human RCC system. Co-culture of A498 cells expressing DsRed-labeled mitochondria (Supplementary 3B) with HUVECs demonstrated DsRed^+^ signal within CD31^+^ endothelial cells by microscopy (Fig. 4F) and ratio-dependent acquisition of DsRed signal by flow cytometry (Fig. 4G). Consistent with these experimental observations, tumor-specific mtDNA mutations were also detected within CD31-laser micro-dissected endothelial compartments from RCC patient specimens (Fig. 4H).

Finally, mitochondrial transfer to endothelial cells was reduced by cytochalasin B, consistent with involvement of actin-dependent processes (Fig. 4I), and was decreased by inhibition of connexin-43 signaling using GAP26^31^ (Fig. 4J). Importantly, cytochalasin B at the concentration used - 400 nM - caused only modest toxicity and did not prevent A498 cell adherence (Supplementary Fig. 3C), suggesting that reduced endothelial acquisition of tumor mitochondria is not attributable solely to gross cytotoxicity.

Together, these results demonstrate tumor-to-endothelium mitochondrial transfer across murine and human RCC models and support the presence of tumor-associated mtDNA mutations within endothelial compartments in human RCC.

### 5. Human TAMAs-specific CD8^+^ T cells recognize endothelial cells presenting mitochondrial neoantigens

To determine the relevance of TAMAs in human cancer, we first assessed the prevalence of tumor-associated missense mtDNA mutations in patient cohorts and examined whether mtDNA mutation status associates with vascular features in RCC^32^. In two independent cohorts comprising 56 primary renal cell carcinomas and 133 breast and ovarian cancers^33^, we found that 33.9% of RCC patients and 30.8% of breast/ovarian cancer patients harbored mitochondrial missense mutations present at ≥30% heteroplasmy (Table 1). Matched adjacent healthy tissue was sequenced when available to validate the tumor specificity of these variants. In RCC cases with paired tumor and adjacent normal tissue available for orthogonal validation, targeted mtDNA sequencing supported the robustness of mutation detection in tumor samples.

**Table 1.**
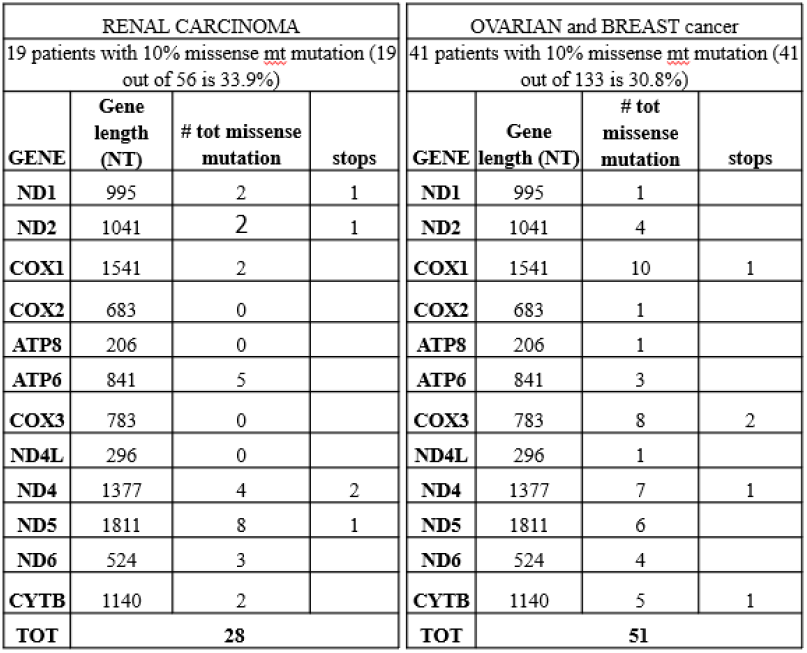
Cancer patients harbor mt missense mutations. Summary of NGS data results from two cohort of cancer patients (56 kidney cancer and 133 breast and ovarian cancer). Respectively, 33.9% and 30.8% of patients harbored at least the 10% of heteroplasmy for mt missense mutation.

We next asked whether mtDNA mutation burden correlated with clinicopathologic parameters in RCC. While no significant associations were observed with renal vein invasion, T stage, or histologic subtype, mutation burden showed a significant association with tumor grade (Supplementary Table 1). We next explored whether mtDNA mutation status associates with immune or vascular staining features in patient RCC samples. In an independent analysis of RCC tumors integrating mtDNA mutation profiling with quantitative immunostaining endpoints, neither the mitochondrial mutational load score nor the total number of mtDNA mutations showed significant associations with the abundance of CD3, CD8, FOXP3, or CD31 staining (Supplementary Table 2). However, when individual recurrent mitochondrial gene mutations were evaluated, the presence of mutations on COX1 were associated with differences in CD31 staining (Fig. 5 A). These findings support the concept that specific mitochondrially encoded alterations, not simply overall mutation burden, may associate with vascular phenotypes in human RCC and provide a clinical rationale for investigating mitochondrial neoantigens in the context of tumor endothelium. In exploratory analyses of progression-free survival (PFS), higher mtDNA mutation burden showed an association with PFS in a multivariable Cox model adjusting for grade and stage; however, interpretation is limited by the very small number of patients in the highest-burden category (Supplementary Fig. 5).

**Figure 5.**
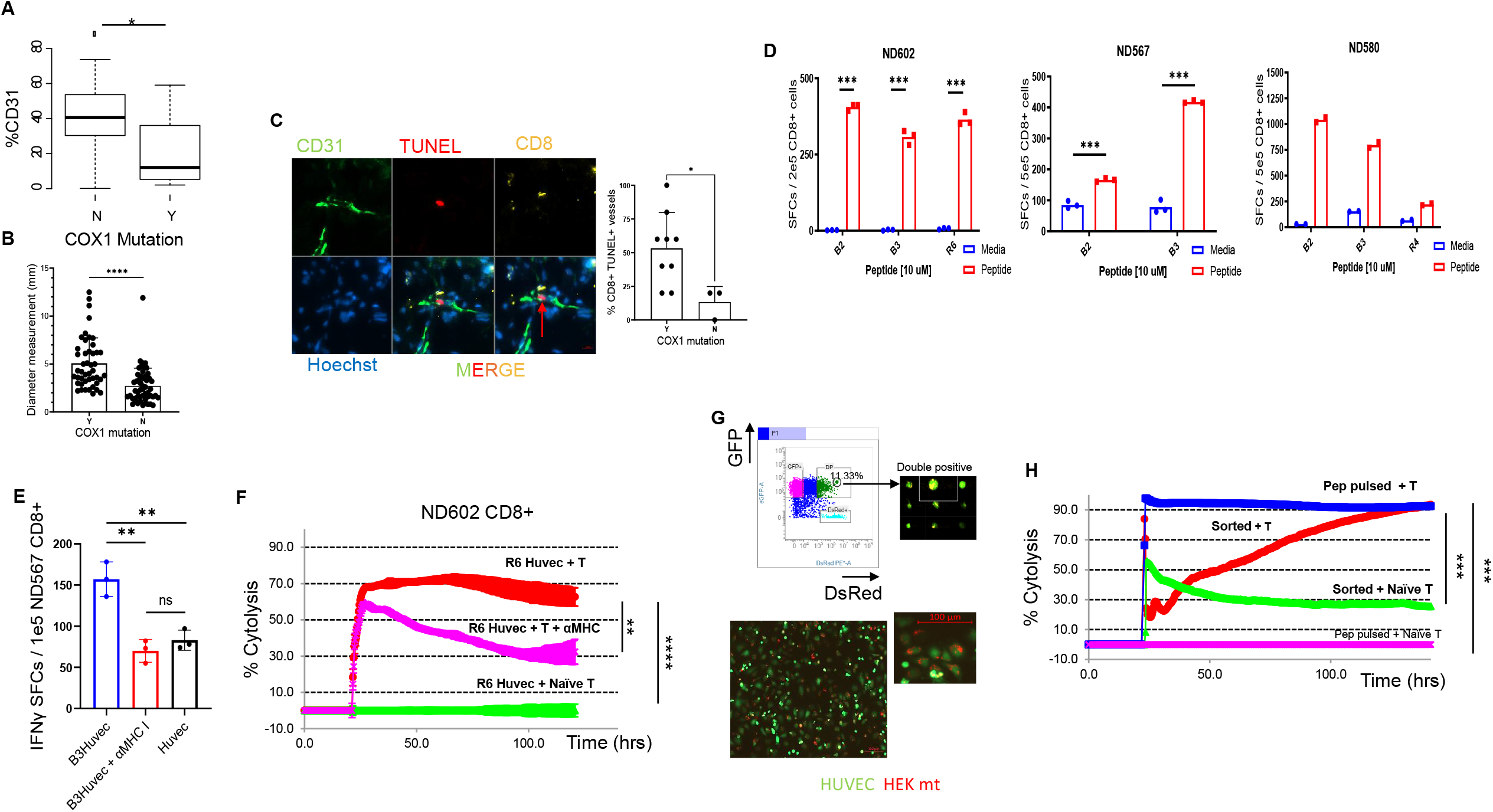
Human relevance: COX1-mutant RCC vascular phenotype and endothelial targeting by TAMA-specific CD8^+^ T cells. (A) CD31 staining levels in RCC samples stratified by presence or absence of COX1 mutation. (B) Quantification of vessel diameter (mm) stratified by COX1 mutation status. (C) Representative multiplex immunofluorescence staining (CD31, TUNEL, CD8) with quantification of CD8-associated apoptotic vessels in COX1-mutant vs COX1–wildtype tumors. (D) IFN_γ_ ELISpot responses of human CD8^+^ T cells expanded against mitochondrial neoantigen peptides, showing responses to peptide stimulation compared with media control. (E) Endothelial cell stimulation assays showing IFN_γ_ production by TAMA-specific CD8^+^ T cells following co-culture with peptide-pulsed HUVECs, with and without MHC class I blockade. (F) Real-time cytotoxicity assay showing killing kinetics of HUVEC targets by TAMA-specific CD8^+^ T cells, with or without MHC class I blockade. (G) Sorting strategy and representative imaging confirming enrichment of endothelial recipient populations following mitochondrial transfer co-culture. (H) Cytotoxicity kinetics using sorted recipient endothelial populations following mitochondrial transfer. Each dot represents one patient sample (A–C) or independent replicate (D–H).

Consistent with a mutation-linked vascular phenotype, stratification of tumors based on COX1 mutation status revealed significant differences in vessel morphology, with COX1-mutant RCCs exhibiting increased vessel diameters compared with tumors in which no COX1 mutation was detected (Fig. 5B). In parallel, multiplex immunofluorescence staining of RCC patient tumors demonstrated the presence of TUNEL^+^CD31^+^ vascular structures in close spatial proximity to CD8^+^ T cells, and quantification showed a significantly higher frequency of CD8-associated apoptotic vessels in COX1-mutant tumors compared with tumors lacking a detectable COX1 mutation (Fig. 5C). Together, these analyses provide human in situ evidence consistent with CD8-associated endothelial injury and further support the clinical relevance of vascular features linked to mtDNA mutation status.

Having established the prevalence of mitochondrial missense mutations in human cancer and identified vascular associations in RCC, we next tested whether these tumor-associated variants can generate immunogenic epitopes recognized by human CD8^+^ T cells. Candidate peptides derived from tumor-specific mitochondrial missense mutations were evaluated for predicted HLA-A*02:01 binding affinity using the publicly available NetMHC4.0 prediction tool^34^. Predicted 9-mer peptides with binding affinities <200 nM were prioritized as strong binders, and four high-confidence candidates derived from recurrent mutations in mitochondrially encoded genes, including COX1 and ND5, were selected for functional validation (Table 2). Using these peptides, we expanded autologous or HLA-matched TAMAs-specific CD8^+^ T cells from multiple healthy donors. In IFN_γ_ ELISPOT assays, expanded TAMAs-specific CD8^+^ T cells mounted strong responses to mutant peptide–pulsed dendritic cells (Fig. 5D), confirming that tumor-associated mitochondrial missense mutations can elicit mutation-directed CD8^+^ T cell reactivity in humans.

**Table 2.**
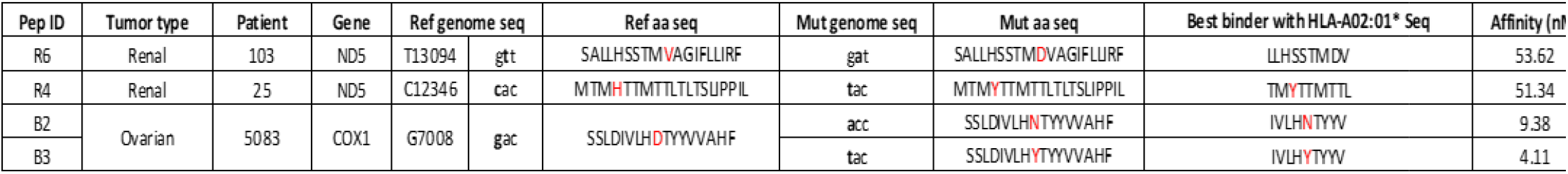
Immunogenic peptide candidates derived from mtDNA mutations of cancer patients. Derivation of the 4 peptides utilized during *in vitro* healthy donor cell priming experiments and their binding affinity score for HLA-A02:01* complex.

We next asked whether human endothelial cells can present TAMAs-derived peptides and serve as direct targets for TAMAs-specific CD8^+^ T cells. Because endothelial antigen presentation is strongly enhanced by inflammatory cues, we stimulated HLA-matched primary human umbilical vein endothelial cells (HUVECs) with IFN_γ_ to upregulate HLA class I expression and antigen presentation pathways (Sup. Fig. 4A-B)^35^. Mutant peptide–pulsed HUVECs induced robust IFN_γ_ responses from TAMAs-specific CD8^+^ T cells, while MHC class I blockade strongly reduced these responses (Fig. 5E), demonstrating that endothelial recognition occurs through classical HLA-I–restricted antigen presentation. Consistent with functional target engagement, TAMAs-specific CD8^+^ T cells efficiently killed peptide-pulsed HUVECs in a real-time cytotoxicity assay, whereas naïve CD8^+^ T cells exhibited minimal killing (Fig. 5F). MHC class I blockade significantly reduced endothelial cytotoxicity (Fig. 5F), further supporting antigen-specific, HLA-I–dependent endothelial targeting.

To establish a physiologically relevant route of antigen acquisition by endothelial cells, we first screened multiple human cancer cell lines for tumor-associated mitochondrial missense mutations and prioritized variants predicted to generate HLA-A*02:01–binding mtDNA-encoded neoantigens (Table 3). This analysis identified several candidate missense variants with predicted immunogenic potential, including a mutation shared between human embryonic kidney (HEK) cells and the Mel624 melanoma cell line, enabling standardized testing of TAMA-specific reactivity.

**Table 3.**
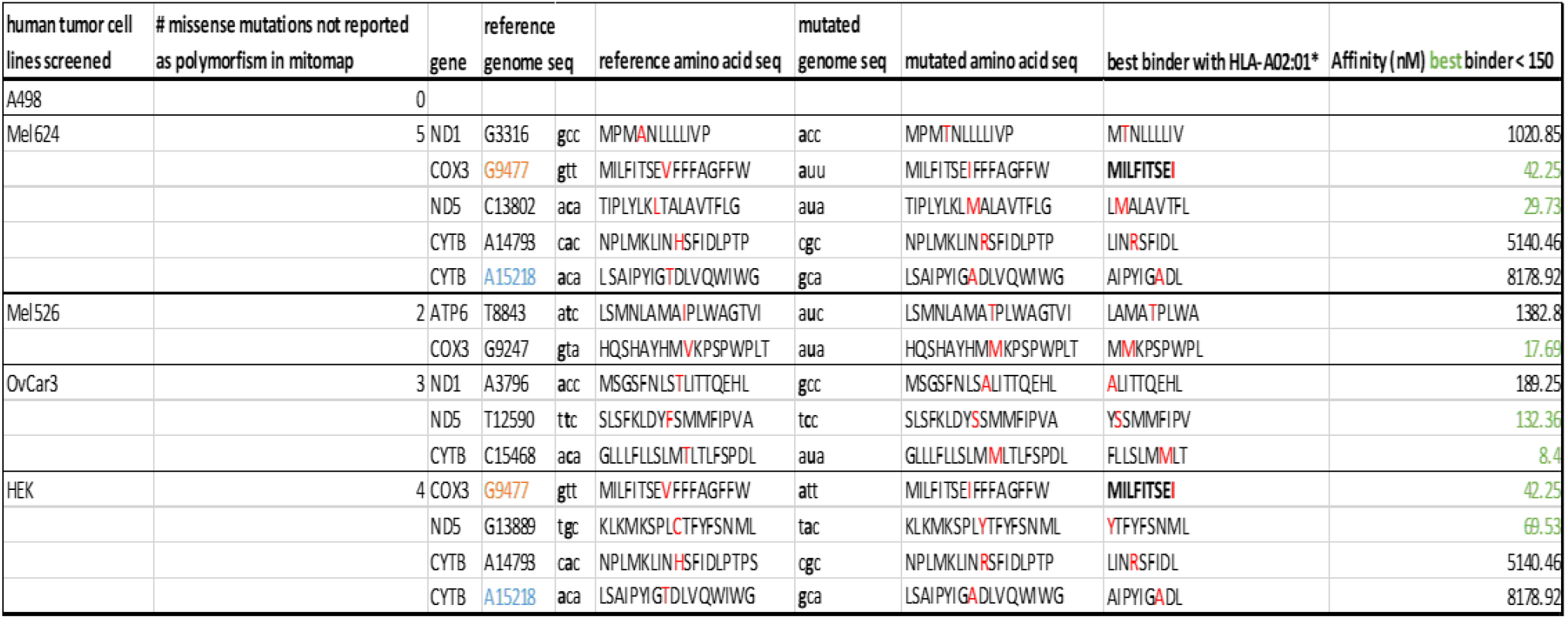
Cancer human cell lines harbor mtDNA missense mutations. Sanger seq data analysis from 5 human cancer cell lines mtDNA, analyzed with Snapgene software: 4 out of 5 analyzed cell lines harbor at least 2 mt missense mutations. 2 mt missense mutations are shared between Mel624 and HEK cell lines. MILFITSEI 9mer has been selected for *in vitro* healthy donor cell priming due to its best affinity with HLA-A02:01* complex.

Building on these findings, we next established an *in vitro* tumor-to-endothelium mitochondrial transfer system in which endothelial cells acquire tumor mitochondria containing mtDNA-encoded neoantigens. We co-cultured immortalized HEK cells harboring the selected mitochondrial missense mutation and engineered to express mitochondria-targeted DsRed with GFP-expressing HUVECs, enabling direct visualization and quantification of mitochondrial transfer. Both microscopy and flow cytometry demonstrated efficient, ratio-dependent acquisition of DsRed-labeled mitochondria by endothelial cells (Fig. 5G and Supplementary Fig. 5B).

Finally, to test whether endothelial recipients of tumor mitochondria can become targets of TAMAs-specific immunity, HEK-DsRed-mito cells were co-cultured with GFP-HUVECs, and endothelial cells were subsequently enriched by CD31 selection and FACS-sorted for GFP + DsRed positivity to eliminate contaminating tumor cells and endothelial cells without HEK derived mitochondria. Sorted endothelial recipient populations were then stimulated with IFN_γ_ to optimize HLA-I presentation prior to coculture with TAMAs-specific CD8^+^ T cells (Supplementary Fig. 3). Under these conditions, TAMAs-specific CD8^+^ T cells efficiently killed endothelial recipient populations following mitochondrial transfer, whereas naïve T cells exhibited minimal cytotoxicity (Fig. 5H). These findings demonstrate that endothelial cells exposed to tumor mitochondrial transfer can become direct targets of antigen-experienced, mitochondrial neoantigen–specific CD8^+^ T cells.

Together, these data establish that (i) tumor-associated mitochondrial missense mutations are prevalent across human cancers and can be tumor-restricted; (ii) in RCC patient samples, specific mitochondrial gene mutations, including COX1, associate with altered endothelial markers consistent with vascular differences and COX1-mutant tumors exhibit in situ signatures consistent with CD8-associated endothelial injury; and (iii) human endothelial cells can present mitochondrial neoantigens via MHC class I and undergo antigen-specific cytotoxic killing by TAMAs-specific CD8^+^ T cells. Collectively, these results support a model in which mitochondrial neoantigens may broaden the immune target landscape beyond malignant cells to include the tumor vasculature.

## Discussion

In this study, we establish a mechanistic framework linking mitochondrial neoantigen immunity to immune-driven remodeling of the tumor vasculature in RCC. We show that vaccination with tumor-associated mitochondrial antigens (TAMAs) elicits antitumor activity in the checkpoint-insensitive RENCA model and that combining TAMAs vaccination with PD-1/PD-L1 blockade further improves tumor control and survival.^15,16^ This therapeutic benefit is accompanied by increased intratumoral CD8^+^ T-cell abundance, enhanced cytotoxic effector features, and reduced accumulation of immunosuppressive myeloid populations. In parallel, TAMAs + ICI promotes vascular remodeling characterized by increased pericyte coverage, reduced vascular leakage, improved perfusion, and diminished hypoxia, features consistent with a more functional and stabilized vascular state.^24,36^ Mechanistically, these vascular changes require CD8^+^ T cells and IFN_γ_ and are associated with increased endothelial apoptosis and reduced CD31 signal. Finally, we demonstrate tumor-derived mitochondria transfer into endothelial cells *in vitro* and *in vivo*, that tumor-associated mtDNA mutations can be detected in endothelial-enriched compartments in murine and human RCC, and that human endothelial cells can present mitochondrial neoantigens and become direct targets of TAMAs-specific CD8^+^ T cells. Together, these findings support a model in which mitochondrial neoantigen immunity enhances tumor cell killing while reshaping the vascular microenvironment in ways that reinforce responsiveness to checkpoint blockade.^27,36^

A central limitation of immune checkpoint blockade in RCC is that durable tumor regression often depends on the presence of sufficiently primed tumor-reactive T cells^37^. In this context, antigen-directed vaccination strategies capable of expanding tumor-specific CD8^+^ T-cell populations represent a rational complement to checkpoint therapy, particularly given the limited availability of highly immunogenic neoantigens in many cancers.^1^ Consistent with this, TAMAs vaccination alone significantly delayed RENCA tumor progression, whereas checkpoint blockade alone showed limited activity. When combined with ICI, TAMAs vaccination increased tumor-infiltrating CD8^+^ T cells and amplified cytotoxic effector signatures. Across tumors, higher intratumoral CD8^+^ T-cell abundance was associated with smaller tumor size and increased tumor apoptosis, supporting a functional contribution of cytotoxic T-cell activity to tumor injury.

Beyond effector differentiation, effective tumor control requires immune access to the tumor parenchyma, a process strongly shaped by vascular architecture and function^27,38^. Tumor blood vessels can limit immune trafficking through heterogeneous perfusion, pathological permeability, hypoxia, and insufficient pericyte stabilization.^36^ Here, the increased pericyte coverage, the reduction in leakage, the coordinated increase in perfusion, and the decrease in hypoxia observed with TAMAs + ICI support an active vascular remodeling program rather than a passive consequence of reduced tumor burden, as perfusion did not correlate with tumor size. These findings are particularly relevant in RCC, where abnormal vascular morphology and angiogenic programs are defining features of disease biology. In clear cell RCC, where VHL loss and constitutive HIF signaling are common, tumors can develop a pseudo-hypoxic state characterized by sustained angiogenesis and vascular dysfunction^39^, which may further constrain immune access and therapeutic efficacy.

Mechanistically, our data identify CD8^+^ T cells and IFN_γ_ as key drivers of immune-associated vascular remodeling. Adoptive transfer studies demonstrate that CD8^+^ T cells in the context of checkpoint blockade increase αSMA^+^ pericyte coverage and increase bigger diameter vessel count, and these effects are abrogated by IFN_γ_ neutralization. CD4^+^ T cells promoted pericyte recruitment more modestly and did not significantly alter vessel caliber, suggesting that CD4^+^ and CD8^+^ T cells contribute through distinct functional programs. In TAMAs + ICI tumors, endothelial apoptosis was increased, and the frequency of TUNEL^+^CD31^+^ vascular structures correlated with intratumoral CD8^+^ T-cell abundance. Taken together, these data support a model in which antigen-specific CD8^+^ T-cell immunity can reshape vascular features that ordinarily constrain immune entry, thereby reinforcing local immune pressure and therapeutic activity^40^. Importantly, the coexistence of normalization-like features (improved perfusion, reduced leakiness, increased pericyte coverage, reduced hypoxia) with endothelial apoptosis suggests a remodeling response that may involve selective injury to vulnerable endothelial elements alongside overall improvement in vascular function.^24,36^

A key conceptual advance of this work is the identification of tumor-to-endothelium mitochondrial transfer as a mechanism that links mitochondrial neoantigens to the vascular compartment. Intercellular mitochondrial transfer has most frequently been discussed in tumors in the context of metabolic adaptation and stress tolerance and has often emphasized stromal-to-tumor mitochondrial donation^20,41^. More recently, mitochondrial transfer has also emerged as a mechanism of immune modulation and immune escape within the tumor microenvironment. Notably, tumor-to–CD8^+^ T-cell transfer of mitochondria carrying mtDNA mutations has been shown to impair T-cell mitochondrial integrity and effector function and contribute to resistance to PD-1 blockade.^42^ In parallel, tumor cell acquisition of mitochondria from immune cells has been reported to promote immune dysfunction and engage tumor-intrinsic inflammatory signaling programs that favor immune evasion.^43^ In contrast to these immune-suppressive paradigms, our findings support a distinct immunological consequence of mitochondrial transfer: tumor-to-endothelium transfer can deliver tumor-derived mtDNA variants to the vasculature, creating an “antigen bridge” that broadens cytotoxic target space beyond malignant cells.

In support of this concept, we demonstrate that mitochondrial missense mutations in human cancers generate immunogenic epitopes capable of activating human CD8^+^ T cells. We used healthy-donor–derived T cells as a proof-of-principle system to validate the immunogenicity and antigen specificity of mitochondrial neoantigen candidates under controlled conditions. Furthermore, human endothelial cells presented TAMAs peptides and activated TAMAs-specific CD8^+^ T cells through classical HLA class I restriction, and endothelial killing was reduced by MHC class I blockade. While endothelial cells are not professional antigen-presenting cells, inflammatory licensing signals such as IFN_γ_ can enhance endothelial HLA class I expression and antigen processing, and checkpoint blockade may further support effector function in this setting^35,44^. Importantly, endothelial recipient populations that acquired tumor mitochondria through co-culture became direct targets of TAMAs-specific CD8^+^ T-cell cytotoxicity after enrichment and sorting, providing functional evidence that mitochondrial transfer can render stromal compartments immunologically visible.

Integrating these observations, we propose a model in which TAMAs vaccination primes a mitochondrial neoantigen–specific CD8^+^ T-cell response that is amplified by checkpoint blockade, leading to both direct tumor cell killing and immune-mediated remodeling of the tumor vasculature. Tumor-to-endothelium mitochondrial transfer may contribute to this process by introducing tumor-derived mtDNA variants into endothelial cells, enabling mitochondrial neoantigen presentation by the vascular compartment and increasing susceptibility of tumor vessels to CD8^+^ T-cell attack. In this framework, endothelial targeting may represent a mechanism by which antigen-specific immunity reshapes the vascular barrier that ordinarily constrains immune infiltration, thereby promoting conditions favorable for immune trafficking and effector function^1,24,45^.

This study also raises important questions for future work. While our enrichment and sorting strategy minimizes tumor contamination, additional routes of endothelial antigen acquisition— such as uptake of mitochondrial debris, extracellular vesicle–associated material, or other intercellular transfer processes, may contribute to antigen loading onto endothelial HLA class I^42,43^. Moreover, mitochondrial neoantigens are expected to be patient- and tumor-specific and may vary by heteroplasmy level, HLA genotype, and inflammatory context, highlighting the need for expanded analyses across diverse RCC cohorts and additional tumor types^45^. In exploratory analyses of progression-free survival (PFS), higher mtDNA mutation burden showed an association with PFS in a multivariable Cox model adjusting for grade and stage; however, interpretation is limited by the very small number of patients in the highest-burden category. Finally, as endothelial targeting could in principle carry risks of off-tumor vascular injury, future studies will be needed to define the therapeutic window, the spatial restriction of endothelial antigen presentation within tumors, and the requirements for selective tumor-associated vascular remodeling without systemic toxicity.

In summary, our findings identify mitochondrial neoantigens as a tractable class of targets and establish tumor endothelium as an immunologically relevant stromal compartment capable of acquiring tumor-derived mitochondrial material. These results suggest that combining mitochondrial neoantigen priming strategies with immune checkpoint blockade may provide a rational approach to enhance antitumor immunity in RCC, in part by improving immune access through vascular remodeling. More broadly, the observation that tumor-derived mitochondria can transfer into endothelium and introduce tumor-associated mtDNA mutations into the vascular compartment raises the possibility that mitochondrial transfer expands the anatomical range of immune targets beyond malignant cells, with therapeutic implications for improving response to immunotherapy in solid tumors.

## Methods

### Mice and cell lines

BALB/c (H-2d) mice (8-wk-old) were purchased from The Jackson Laboratory. Murine renal carcinoma RENCA cell line, human kidney cancer A498, Caki-1, 786-O, Mel624, Mel526, OvCar3, HUVEC, HEK-293 and endothelial MS1 cell line were purchased from the American Type Culture Collection. BALB/c Mouse Primary Lung Microvascular Endothelial Cells were purchased from Cellbiologics. Murine mesothelioma AB12 cell line was purchased from Millipore Sigma. HLA-A*02:01 healthy donor cells were purchased from Human Immunology Core at Perelman School of Medicine, University of Pennsylvania. Beatriz M. Carreno, University of Pennsylvania, donated CD40L-expressing K562 cells. GFP HUVEC (Product Code: ZHC-2402) were purchased from Cellworks.

### Generation of mtGFP -labelled RENCA cells

mtGFP-labelled RENCA cells were produced by transfecting RENCA cells with the plasmid PEGFP-N1-mt-ro1GFP #82407 purchased from Addgene. This plasmid contains a cytomegalovirus (CMV) promoter-driven ro1GFP reporter gene, which is fused to the leader sequence of the E1α subunit of pyruvate dehydrogenase for mitochondrial targeting (N terminal on insert). 2.5 □ µg of DNA for 0.3 □ × □ 10^6^ cells was used for transfection with Lipofectamine 3000, following the manufacturer’s protocol (Thermo Fisher Scientific; #L3000001). Two days after transfection, cloning is 96 well plates was started and after 10 days 100 µL of medium were added to the wells that presented 100% of GFP cells under fluorescence microscopy.

### Generation of mtDsRed-labelled A498 cells and HEK-293 cells

mtDsRed-labelled A498 cells or HEK-293 cells were produced by transfecting A498 cells or HEK-293 with the plasmid pLV-mitoDsRed #44386 purchased from Addgene. This plasmid contains a cytomegalovirus (CMV) promoter-driven mitoDsRed reporter gene, which is fused to a 61 aa targeting sequence of the P1 isoform F1F0-ATP synthase (N terminal on insert). 2.5 □ µg of DNA for 0.3 □ × □ 10^6^ cells were used. Two days after transfection, cloning in 96 well plates was started and after 10 days 100 □ µL of medium were added to the wells that presented 100% of DsRed cells under fluorescence microscopy.

### Immunohistochemistry and Immunofluorescence

Tumors or healthy tissues were fixed in PFA 3.7 %+ sucrose 2% for one hour at room temperature on the wheel, then 30 minutes with just sucrose 2% and following incubation at 4°C in sucrose 20% until tissue reach the bottom of the Eppendorf. The day after, tissue were embedded in sucrose 20% and OCT (Scigen #23-730-625) with a 1:1 ratio until tissue reaches the bottom of the mold. Finally, tissues were embedded in OCT and immediately snap-frozen in dry ice. Tissue OCT embedded were sectioned in 7 µm thick sections with a Leica CM1950 cryostat (Leica, Wetzlar, Germany) and stained for CD8 (1:300, Cell Signaling, #98941) with standard procedure for immunohistochemistry and manufacturer’s instructions by the Penn Vet Comparative Pathology Core (CPC) at University of Pennsylvania, which is partially supported by the Abramson Cancer Center Support Grant (P30 CA016520). The Aperio Versa 200 scanner used for whole slide imaging and the image analysis software was supported by an NIH Shared Instrumentation Grant (S10 OD023465-01A1). The Leica BOND RXm instrument used for IHC was acquired through the Penn Vet IIZD Core pilot grant opportunity 2022. For immunofluorescence (IF) analysis, the sections or the cells were fixed (if not previously done) in ascending concentrations of ethanol, blocked with 8% bovine serum albumin for 1 hour and incubated with the primary antibody PD-L1 (1:100, ab80276, Abcam, Cambridge, UK), CD31 (1:200, Pharmingen # 550274), CD31 (1:50, ab28364, Abcam, Cambridge, UK), NG2 (1:50, Thermofisher, #MA5-24247), α-Smooth Muscle – Cy3 (1:50, C6198, SIGMA), CD8 (1:200, ab22378, Abcam, Cambridge, UK), CD31(1:20, 77699T, Cell Signaling), CD8 (1:800, 66868-1, Proteintech). Slides were then washed and incubated with the labelled Alexa Fluor 594 goat anti-rat IgG (H+L) (Invitrogen, A11007) diluted 1:400 or Alexa Fluor 488 goat anti-rabbit or anti-rat IgG (H+L) (Invitrogen, A11006) diluted 1:800. Finally, nuclei were stained with Hoechst 10 μg/mL (invitrogen, H3570) and fixed with 2% paraformaldehyde. To analyze the levels of apoptosis in the TME, tumor tissues were stained using an In Situ Cell Death Detection kit (Roche, 12156792910, TUNEL) according to the manufacturer’s instructions. Slides were ounted with Fluoromount-G™ Mounting Medium (Invitrogen, 00-4958-02) and steal with nail polish. For each specimen, random pictures or whole slide scan were collected with a Zeiss Observer.Z1 inverted Fluorescent microscope with an AxioCam MRm camera (Zeiss). Quantitative image analysis was performed using FIJI open source software or Zen 3.3 software. Measurements of tumor vessels diameter were calculated with FIJI using the plugin “Diameter Measurements”. Slides were analyzed in a blinded fashion.

### RT-PCR

The relative quantification of the expression levels of selected genes was carried out by qPCR. Total RNA from cells and tissues was extracted using TRIzol reagent (Invitrogen, 15596018) according to the manufacturer’s instructions. Reverse transcription and RT-PCR reactions were carried out using the High Capacity cDNA Reverse Transcription Kit (Thermo Fisher Scientific, 4368814) and TaqMan Gene Expression Master Mix (Thermo Fisher Scientific, 4369016) according to the manufacturer’s instructions. Runs were performed using the QuantStudio 6 Flex Real-Time PCR System (Thermo Fisher Scientific). All TaqMan primers were purchased from Thermo Fisher Scientific. Housekeeping gene Rn18s (Mm03928990_g1), PDL-1 (Mm03048248_m1), Perforin (Mm00812512_m1), Ang1 (Mm01233546_g1), Ang2 (Mm00657574_s1), VECAD/Cdh5 (Mm00486938_m1), Vcam1 (Mm01320970_m1), Cxcl10 (Mm00445235_m1), Cxcl11 (Mm00444662_m1), Cxcl9 (Mm00434946_m1), Vegfa (Mm00437306_m1), Pdgfb (Mm00440677_m1), Irf3 (Mm00516784_m1), CD8a (Mm01182107_g1), Ifnγ (Mm01168134_m1), Il12a (Mm00434169_m1), Il12b (Mm01288989_m1), and Gzmb (Mm00442837_m1), TNF (Mm00443258_m1), PTEN (Mm00477208_m1), Itgax (Mm00498701_m1), Tnfsf10 (Mm01283606_m1), Tnfrsf1a (Mm00441883_g1).

### Murine BMDCs generation and maturation

Bone marrow cells were isolated from hind leg femurs and tibias of mice, and DCs were isolated, cultured, and matured as previously described^46^.

### BMDCs-based immunotherapy

Mitochondria from RENCA solid tumors and RENCA cell lines, as well as control mitochondria, were purified, co-cultured with BALB/c-derived DCs and matured as described above. In the therapeutic protocol, mice were injected s.c. with 10^6^ RENCA cells and, starting 1 week later, they were given three injections of 10^6^ DCs at weekly intervals. For some mice, αPDL-1 (BioXcell; clone 10F.9G2, BE0101) or αPD-1 (BioXcell; clone RMP1-14, BE0146) was also administrated every 3 days starting from the day before the first vaccination, 200 □ µg/mouse^3^. Tumor progression was monitored every other day from the tumor challenge.

### Synthetic peptides

Costume peptides COX1291–306 (MFTVGLDVDTRTYFT), COX1295–310 (GLDVDTRTYFTSATM), COX1299–314 (DTRTYFTSATMIIAI), B2 (IVLHNTYYV), B3 (IVLHYTYYV), R4 (TMYTTMTTL) and R6 (LLHSSTMDV) were synthesized by and purchased from Mimotopes. Lyophilized peptides were dissolved in 10% DMSO (Mylan Cryoserv 67457-178-50) in sterile water pH 7.4 and passed through a 0.2 □ uM Centrex filter (10467013).

### Purification of mitochondria and mtDNA extraction

Mitochondria were purified from RENCA cell lines or tissues (RENCA solid tumor and kidney) using the discontinuous sucrose gradient method described previously. Preparations were frozen (liquid nitrogen) and thawed (37°C, water bath) for three cycles, sonicated, and irradiated at 5000 rad for sterilization. Quantification was carried out with the BCA assay. mtDNA was isolated from cell lines using a Mitochondrial DNA Isolation Kit from BIOVISION.

### Murine IFN-_γ_ ELISPOT assay

Isolated CD3+ T cells from the spleen of tumor-bearing mice were co-cultured for 16 h with DCs pulsed with RENCA mt lysate or COX1 peptides pool at a 10:1 ratio, if not differently specified. After incubation, plates were washed with PBS and Tween 20, ELISPOT was performed and spots were counted using an automated ELISPOT reader (Autoimmun Diagnostika GmbH)^47^.

### Intracellular cytokine staining for IFN-_γ_

A total of 10^6^ mouse splenocytes was incubated with the indicated peptides (final concentration of each peptide, 10 mg/ml) and 1 mg/ml Golgi Stop (BD Biosciences, Pharmingen) at 37°C for 12–16 h and labeled with CD3+/CD4+ or CD8+ and IFN-_γ_, as previously described^48^. Cells were analyzed on a FACSCanto flow cytometer using FlowJo software.

### Flow cytometry

Cells or processed tissues were labeled with Abs from BioLegend (VE-cadherin/CD144 [clone BV13], CD45 [clone S18009F], CD3 [clone 17A2], CD8a [clone 53-6.7], CD4 [gk1.5], PD-L1 [clone 10F.9G2], CD11b [clone M1/70], CD86 [clonGL-1], F4/80 [clone BM8], CD11c [clone N418], IFN_γ_ [clone XMG1.2], GR1 [clone RB6-8C5], FoxP3 [clone NRRF-30], CD62L [cloneMEL-14], PD1 [clone 29F.1A12], CD25 [clone 3C7] TNF_α_ [cloneMP6-XT22], CD107 [clone 1D4B], CD31 [clone 390]), from abcam (NG2 [clone EPR23752-147]), from BD Pharmingen (anti-Human CD31 [clone M89D3], VEGFR2 [clone Avas 12 α1]), from eBioscience (anti-Human CD11c, [clone 3.9], anti-Human CD11c CD83 [clone HB15e], anti-Human CD11c CD86 [clone IT2.2], anti-Human HLA-ABC [clone W6/32], anti-Human HLA-A2 [clone BB7.2]). Labeled cells were evaluated with a FACSCanto and the data were analyzed using FlowJo software (V10.8.1_CL).

### Western blot analysis

Cells were collected in ice-cold PBS and proteins were extracted using 1× RIPA buffer supplemented with protease (Sigma, B8640) and phosphatase inhibitors (Sigma, P5726 and P0044). Tissues were snap frozen and processed in the same way. Proteins were separated by 4–15% SDS–PAGE, transferred to polyvinylidene fluoride (PVDF) membranes and blocked with 5% nonfat milk or BSA in 1× PBS with Tween-20 (0.1%). Primary antibodies against Perforin (Cell Signaling Technology, 31647), Perforin (Cell Signaling Technology, 62550), Granzyme B (Cell Signaling Technology, 4275), CD31 (Cell Signaling Technology, 77699), Vinculin (Cell Signaling Technology, 13901), GFP (Cell Signaling, 2956) and Calreticulin (Cell Signaling, 19780) were added at 1:1,000 and incubated overnight at 4 □ °C. Membranes were washed, and secondary antibodies (ThermoFisher, 31460, RRID: AB_228341 or 31430, RRID: AB_228307) were added at 1:2,000. ECL (Thermo Scientific, 32106 and GE Healthcare, RPN2232) was added, and membranes were exposed to ChemiDoc (Bio-Rad).

### Ultrasound analysis of tumor vasculature

B-mode ultrasound images (VisualSonics, VevoLAZR, 21-MHz linear transducer, Fujifilm, Toronto, Canada) of each RENCA tumor were acquired to assess the tumor size and positioning. This same equipment was used to acquire contrast-enhanced ultrasound (CEUS) scans to assess tumor perfusion; fixed imaging parameters were used (18 MHz; contrast gain = 41; 2D gain = 18; sensitivity = 3; standard line density; power = 4). An intravenous bolus injection of 0.01 mL microbubbles was used for CEUS imaging. Tumor perfusion was assessed using VevoLab software (Fujifilm, VisualSonics, Toronto, Canada). The borders of each tumor were manually outlined in the initial B-mode image to create a region of interest (ROI). The mean intensity of the ROI was fitted to a bolus perfusion model, and the following parameters were assessed: peak enhancement (PE - the difference between the maximum amplitude and baseline intensity, which is proportional to microbubble concentration and indicative of relative blood volume); perfusion index (PI - the area under curve divided by mean transit time of flow); and time to peak (TTP - time from contrast administration to peak enhancement)^49^.

### Evans Blue perfusion assay

Evans Blue (MP Biomedicals, LLC, 151108) was concentrated at 50 mg/mL and injected intravenously to each mouse 0.01 mL per gram of mouse weight. After 30 minutes, mice were euthanized and perfused with 50 mL of 50 mM sodium citrate solution to remove the excess Evans Blue from within the blood vessels. Tumors were collected in a 1.5 mL Eppendorf tube, weighted and dried in a hot plate at 56 □C for 24 hours. 500 μL of formamide were added and tissues were homogenized and incubated at 56 □C for 2 days. Tissues were centrifuged at max speed for 5 minutes to allow Evans Blue collection. absorbance of samples and standards was measured at 620 nm^50,51^

### EF5 Hypoxia Detection Assay

2-nitroimidazole EF5 [2-(2-nitro-1H-imidazol-1-yl)-N-(2,2,3,3,3-pentafluoropropyl)acetamide] (#EF5-30C3, Sigma-Aldrich) is a compound developed at the University of Pennsylvania by Dr. Cameron Koch and Dr. Sydney Evans. Upon injection into animal tissues (tail-vein injection using 10 mM drug with the amount (in mL) equal to 1/100 of the animal weight, in grams), EF5 selectively binds to hypoxic cells and forms adducts. After 2 hours from injection, mice are euthanized and tumor tissues are harvested and embedded in OCT, as described before. Slides of 7μm are fixed with ethanol, blocked with 8% bovine serum albumin (BSA) in PBS-TT (PBS with 0.1% Tween-20 and 0.025% Triton X-100) for 1-2 h at room temperature in humid chamber, or 4h at 4 □C. A mouse monoclonal antibody, clone ELK3-51, which is directly conjugated to Cyanine 3 is then used to selectively bind the EF5 adducts overnight at 4°C. Finally, nuclei are stained with Hoechst 10 μg/mL (invitrogen, H3570) and fixed with 2% paraformaldehyde. Slides were acquired and analyzed as described above.

### Adoptive cell transfer

Tumor-free BALB/c mice were vaccinated three times with TAMAs vaccine or control DCs) and αPDL-1, and sacrificed 1 week later. CD3+, CD4+, and CD8+ T cells were magnetically isolated from the spleens. Isolated T cells were injected i.v. into RENCA tumor–bearing mice (challenged 16 days before transfer). Half mice were treated with αIFNγ (BioXCell, clone R4-6A2, BE0054).

### Computational prediction of neo-epitopes

We investigated the mtDNA of tumor patients to detect immunogenic mutations able to trigger human cytolytic lymphocytes. We collected data from 2 cohorts of patients: NGS data on a cohort of 56 kidney cancer patients and, thanks to the Basser Center at the University of Pennsylvania, Whole exome sequencing (WES) data from a cohort of 133 BRCA carriers. We found respectively, 33.9% and 30.8% of patients harboring at least 10% of heteroplasmy for mt missense mutations. Mitochondria neo-antigens derived from these mutated mtDNA sequences were scored for HLA-A*02:01 binding affinity by the publicly available NetMHC4.0 prediction tool^34^. 9mers with binding affinity prediction effective with a concentration lower than 200 nM were considered as strongest binders and used for healthy donor PBMCs priming in vitro.

### Human DCs generation

DCs were generated by the “adherence method”. Briefly, peripheral blood mononuclear cells (PBMCs) were purchased from Human Immunology Core from University of Pennsylvania. A total of 1 □ × □ 10^8^ mononuclear cells were plated in RPMI containing 2 mM GlutaMAX-I (gibco, #35050-061)+ 10 mM HEPES (gibco, #15630-080) + non-essential amino-acids (gibco, #1140-050) + Penicillin Streptomycin Solution (corning, #30-002-CI) + 1% AB Human sera not inactivated (Sigma-Aldrich, #H3667) in T-75 flasks for 2 hours at 37 □C. The nonadherent cells were removed by washing with phosphate-buffered saline (PBS), and the adherent cells were cultured with 100 ng/mL granulocyte-macrophage colony-stimulating factor (GM-CSF) (Peptrotech, #300-03) and 20 ng/mL interleukin-4 (IL-4) (GensScript, #Z02925-10) for 5 days. IL-4 and GM-CSF were replenished on day 2 and 4. On day 5, cells were harvested and concentrated at 1 ×10^6^/mL with 200 ng/mL GM-CSF, 40 ng/mL IL-4 and, for maturation, 100 U/mL IFN-_γ_ (Preprotech, # 300-02), 5μg/mL poly I:C (InvivoGen, #tlrl-pic) and R848 (InviveGen, #tlrl-r848) with irradiated (10,000 RADS) CD40L-expressing K562 cells. The day after the maturation is confirmed by flow cytometry analysis.

### Human CD8+ T cells priming

Autologous CD8+ T cells were purchased from Human Immunology Core from University of Pennsylvania. Purified CD8+ T cells (5 □ × □ 10^6^ cells/ml) were cultured at a 20:1 ratio with irradiated (2500 Rads) autologous mDC pulsed with peptide (40 □ μg per 1□×□10^6^ DC/ml) in 24-well trays in Optimizer CST media (Gibco) supplemented with 5% pooled human sera. Human IL-7 (10□ng/ml, Preprotech, #200-07), IL-15 (5□ng/ml, Preprotech, #200-15), and IL-12 (10□ng/ml, Preprotech, #200-12) were added on day 0. Fresh media supplemented with IL-7 (10□ng/ml) and IL-15 (5□ng/ml) was added on day 7. Fourteen days after primary mDC stimulation, T cell cultures were harvested and re-stimulated with irradiated (2500 Rads) peptide-pulsed mDC. Cell culture media was supplemented with 50□U/ml IL-2 (Peprotech, #200-02) starting day 2 then every 48□h following secondary stimulation. On days 10–14 of secondary mDC stimulation, antigen-specific T cell responses were identified by IFN-_γ_ ELISPOT assay.

### Human IFN-_γ_ ELISPOT assay

CD8+ T cell reactivity to peptide antigen was assessed by interferon-γ (IFN-_γ_) ELISPOT assay. The spot number was determined in an independent blinded fashion (ZellNet Consulting, New York, NY) using the high-resolution automated KS ELISPOT reader (Zeiss, Thornwood, NY) and KS ELISPOT 4.9.16 software with reading parameters established per International Harmonization Guidelines. A positive response was recorded if the number of spots in the peptide-exposed wells was two times (or more) higher than the number of spots in the unstimulated wells and if there was a minimum of 20 (after subtraction of background spots) peptide-specific spots per 5□×□10^5^ CD8+ cells.

### mtGFP and mitotracker co-localization

To ensure that GFP signal belongs to mitochondria, we used the Mito-Tracker Red CMXRos (Invitrogen, M7512) in order to demonstrate the co-localization of green and red signals. RENCA mtGFP were plated in 24 well plate (at least 20.000 cells) with a round cover glass coated with 0.1% gelatin in PBS. The day after medium was removed and 50nM of Mito-Tracker were added in RPMI for 45 minutes. After 3 PBS washes, Hoechst was added for 10 minutes at 4 □C and the cover glass was mounted on a slide and analyzed at the microscope.

### In vitro mitochondrial transfer by co-culture

Co-culture experiments involved murine MS1 cells, BALB/c mouse primary lung microvascular endothelial cells and mtGFP-labelled RENCA, or human HUVEC, A498, mtDsRed-labelled A498, Caki-1 and 786-O cells on 1% gelatin-coated plates in half ECs medium and half tumor cells medium, for 48 hours if not differently specified. The mitochondrial transfer in murine cells was assessed by co-culturing mtGFP-RENCA cells and MS1 endothelial cells with a 1:2 ratio or endothelial cells isolated from BALB/c mouse lungs with different donor-to-recipient ratios, from 1:1 to 1:5, with total cell not exceeding 3□× □10^4^ per well, if not differently specified. The mitochondrial transfer in human cells was assessed by co-culturing mtDsRed-labelled A498 and HUVEC with different donor-to-recipient ratios from 1:2 to 1:20, with total cell not exceeding 3.5□× □10^6^ per well, if not differently specified. Cytochalasin B from Drechslera dematioidea (Sigma, 14930-96-2) was used at the concentration of 400 nM for human cells coculture to block TNTs formation. Gap26 trifluoroacetate (Sigma, 197250-15-0) was used at the concentration of 175 μM for human cells coculture to test involvement of connexin43 complex to transfer mt by TNTs mechanism^52^.

### Cells sorting

RENCA tumors were collected, stained and sorted using the CD31+, VE-cadherin+ and VEGFR2+ markers. CD31+ and CD31-cells were sorted from renal cell carcinoma tumor cells and HUVEC cells co-culture. All sorting were executed with FACS Aria II, excepted for the GFP+DsRed+ HUVEC that were sorted with BD FACSDiscover™ S8 Spectral Cell Sorter.

### Laser Capture Microdissection

Patients’ blood vessels were stained, isolated and sequenced by Skin Biology and Diseases Resource-based Center at University of Pennsylvania.

### Computational analysis of WES data for mtDNA variant discovery

Whole-exome sequencing (WES) data were obtained as described previously (accession: PRJNA751555)^33^. Reads mapping to the mitochondrial genome were extracted from aligned BAM files and re-aligned to the revised Cambridge Reference Sequence (rCRS; NC_012920.1). Mitochondrial single nucleotide variants (mtSNVs) and small indels were called using the GATK4 mitochondrial short variant discovery pipeline (GATK vX.X). Variant allele fraction (VAF) was used as an estimate of heteroplasmy. Variants were filtered based on mapping quality, base quality, strand bias metrics, and minimum coverage thresholds. Only missense variants meeting filtering criteria and exceeding a predefined heteroplasmy threshold (VAF ≥ X%) were retained for downstream epitope prediction analysis. For BRCA cohort, the raw capture-targeted, WES and RNA sequencing data generated as part of this study have been deposited in the NCBI SRA database under accession code.

### mtDNA amplification and sequencing

Protein-coding sequences were amplified using PCR with 22 pairs of overlapping primers. PCRs were carried out in 25-μl reactions using Phusion DNA polymerase (New England Biolabs) and 10 ng mtDNA template. Prior to sequencing, PCR products were cleaned up using a PureLink Quick PCR Purification Kit (Invitrogen; Life Technologies), and DNA concentration was determined by NanoDrop. Sanger Sequencing was performed on an ABI 3770 (Applied Biosystems) at the Penn Genomics Analysis Core of the University of Pennsylvania. Sequencing was performed with the same primers used for PCR. Mutations were found using SnapGene software.

### HLA PCR and sequencing

Sequences were amplified using PCR with 2 pairs of overlapping primers (12ws/102# with 12ws/209# for HLA-A02:01, and 12ws/01# with 12ws/02# for housekeeping). PCRs were carried out in 25-μl reactions using Phusion DNA polymerase (New England Biolabs) and 260 ng DNA template. Prior to sequencing, PCR products were cleaned up using a PureLink Quick PCR Purification Kit (Invitrogen; Life Technologies), and DNA concentration was determined by NanoDrop. Sanger Sequencing was performed on an ABI 3770 (Applied Biosystems) at the Penn Genomics Analysis Core of the University of Pennsylvania. Sequencing was performed with the same primers used for PCR^53^. Mutations were found using SnapGene software.

### Data analysis

Statistical analyses for the indicated comparisons were performed using the pairwise Student t test. All p values presented are two sided. The log-rank test was performed for the survival curves. Single-step multiple-comparison procedure Tukey test was performed for repeated measurements at different time points. Pearson r test was performed for correlations. All experiments were performed a minimum of three times; in the case of in vivo studies, each group consisted of 10 animals, unless otherwise noted. All figures portray one representative experiment. *p<0.05, **p<0.01, ***p<0.001, n.s. (non-significant).

### Study approval

The Institutional Animal Care and Use Committee and University Laboratory Animal Resources at the University of Pennsylvania approved all animal studies. Mice were treated in accordance with University of Pennsylvania guidelines.

### Cytotoxicity assay

Cytotoxic killing of target cells was assessed using a real-time, impedance-based assay with xCELLigence Real-Time Cell Analyzer System (ACEA Biosciences). Briefly, target cells treated with 10 ng/mL of hIFN_γ_ for 24 hours, pulsed with the immunogenic peptide for 2 hours in the incubator if specified (40□μg per 1□×□10^6^ cells/ml), and then 4 x 10e5 were seeded to the 96-well E-plate. 50 ug/mL of αMHC were added when specified. After 24 hours, 2 x 10e5 CD8+ T cells were added to the wells in 4 : 1 E:T ratio. Tumor killing was monitored every 20 min over 5 days. Significant differences between groups were assessed by two-way ANOVA.

## Notes

### Competing Interest Statement

The authors have declared no competing interest.

### Funding

- NIH: PQ5

