## supplementary figures captions for "“Tumor-to-endothelium mitochondrial transfer licenses endothelial cells for CD8^+^ T cell recognition via mitochondrial neoantigen presentation”"

**Supplementary Figure 1. TAMAs + ICI enhances CD8 activity, apoptosis, and reduces myeloid cells**

(A) Representative flow cytometry plots and quantification of intratumoral CD8⁺IFNγ⁺ T cells in RENCA tumors across treatment groups (DCs, DCs+ICI, TAMAs, TAMAs+ICI). (B) Correlation between intratumoral CD8⁺ T-cell abundance and TUNEL⁺ apoptotic nuclei within tumors. (C) Correlation analyses testing associations between CD4⁺ T-cell infiltration, tumor burden (mm³), and tumor apoptosis. (D) Flow cytometric quantification of intratumoral CD11b⁺Gr1⁺ myeloid cells across treatment groups. Each point represents an individual tumor. Statistical comparisons were performed using one-way ANOVA with multiple-comparison correction.

**Supplementary Figure 2. Tumor perfusion index does not correlate with tumor size**

(A) Peak enhancement / perfusion parameters from contrast-enhanced ultrasound imaging across treatment groups, supporting improved vascular function in treated tumors. (B) Correlation between tumor perfusion index and tumor volume (mm³), showing no significant relationship, consistent with perfusion changes not being solely explained by reduced tumor burden.

**Supplementary Figure 3. Validation of mitochondrial reporters and Cytochalasin B toxicity control**

(A) Representative fluorescence microscopy image of RENCA mtGFP cells confirming mitochondrial reporter localization. (B) Representative fluorescence microscopy image of A498 mitoDsRed cells confirming mitochondrial reporter localization. (C) Brightfield images and cell counts after treatment with vehicle (DMSO) or cytochalasin B at the indicated time points after seeding, supporting that the concentrations used in mitochondrial transfer inhibition experiments caused only modest effects on adherence/viability.

**Supplementary Figure 4. IFNγ licenses endothelial cells for HLA class I presentation**

(A–B) Flow cytometry gating strategy for HUVECs and quantification of HLA class I expression following IFNγ stimulation, supporting endothelial “licensing” prior to peptide-pulsing and coculture assays.

(C) Quantification of mitochondrial transfer frequency in the HEK–HUVEC coculture system at the indicated ratios/conditions (as used for downstream cytotoxicity assays). Each point represents an independent replicate.

**Supplementary Figure 5. Progression-free survival (PFS) by mtDNA mutation burden. Kaplan–Meier curves are shown for mtDNA mutation burden strata.**

Exploratory multivariable analyses suggested a potential association between higher mtDNA mutation burden and PFS, but estimates were unstable due to the small number of high-burden cases.
