## supplementary figures for "“Tumor-to-endothelium mitochondrial transfer licenses endothelial cells for CD8^+^ T cell recognition via mitochondrial neoantigen presentation”"

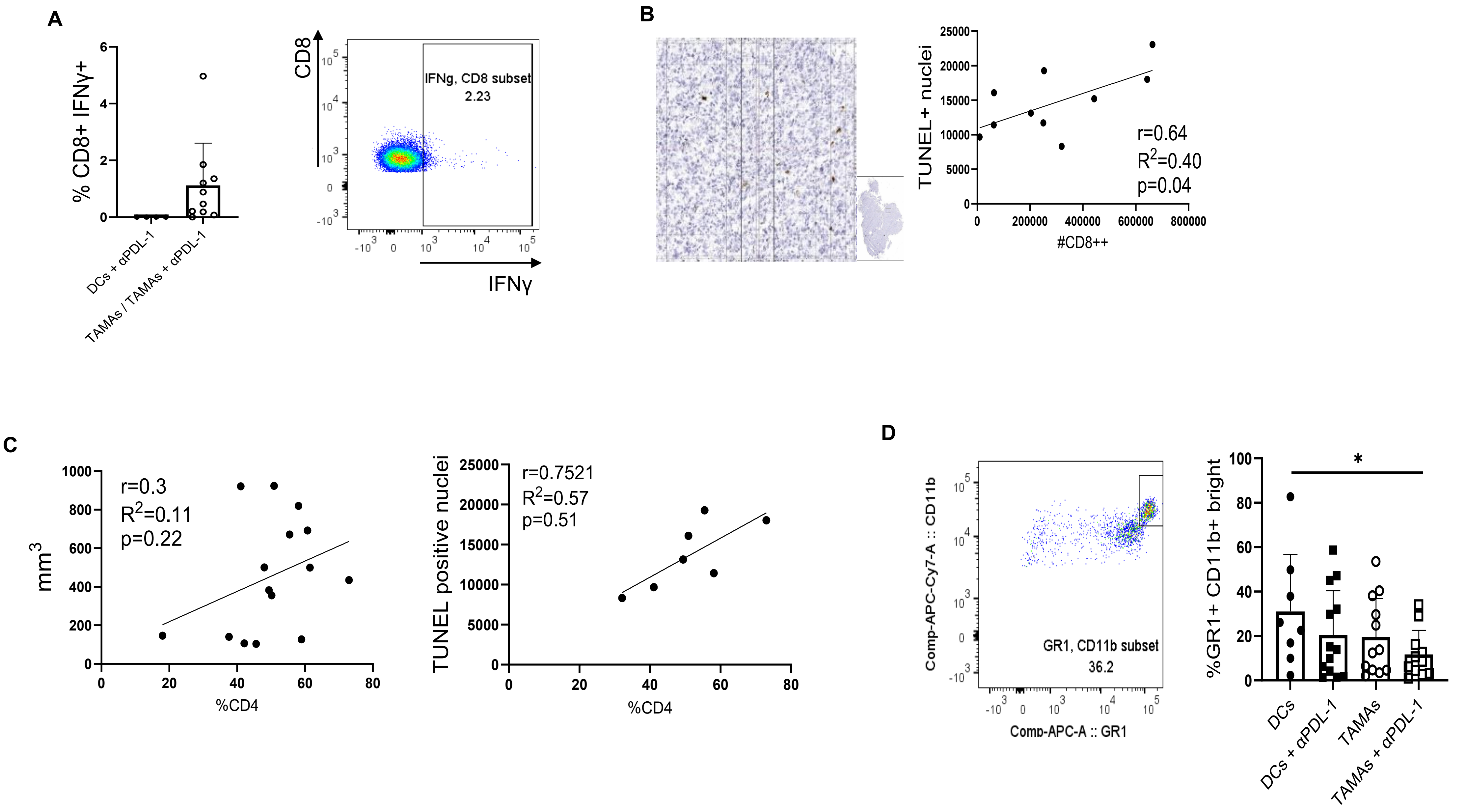


**Supplementary Fig. 1**

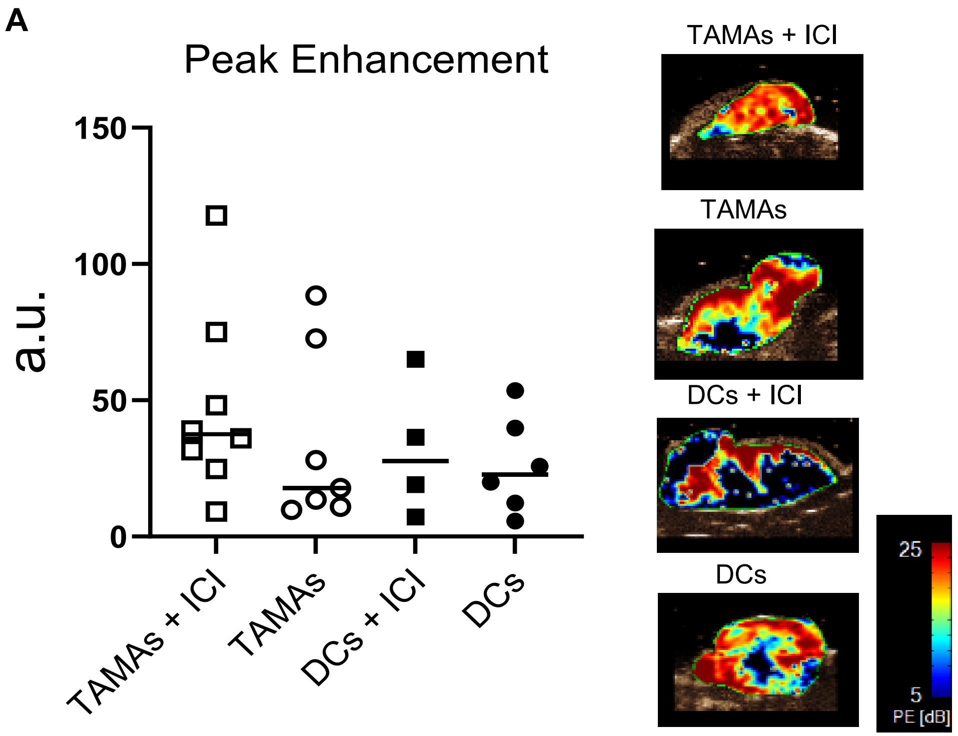


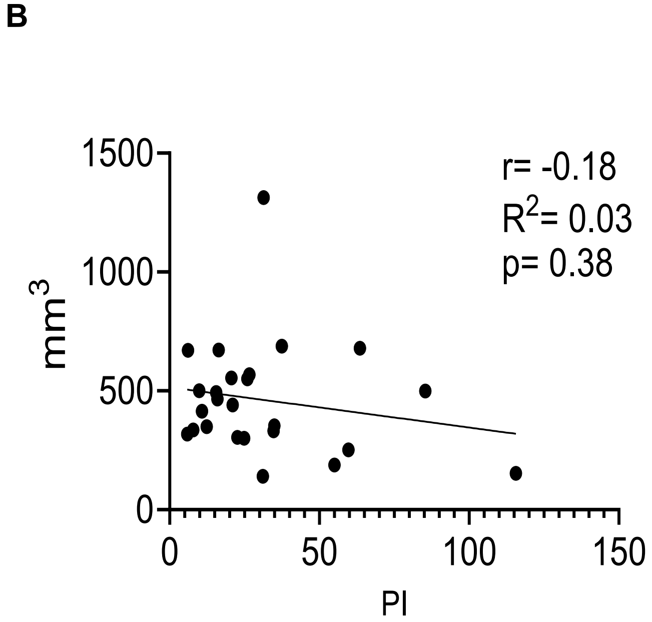


**Supplementary Fig. 2**

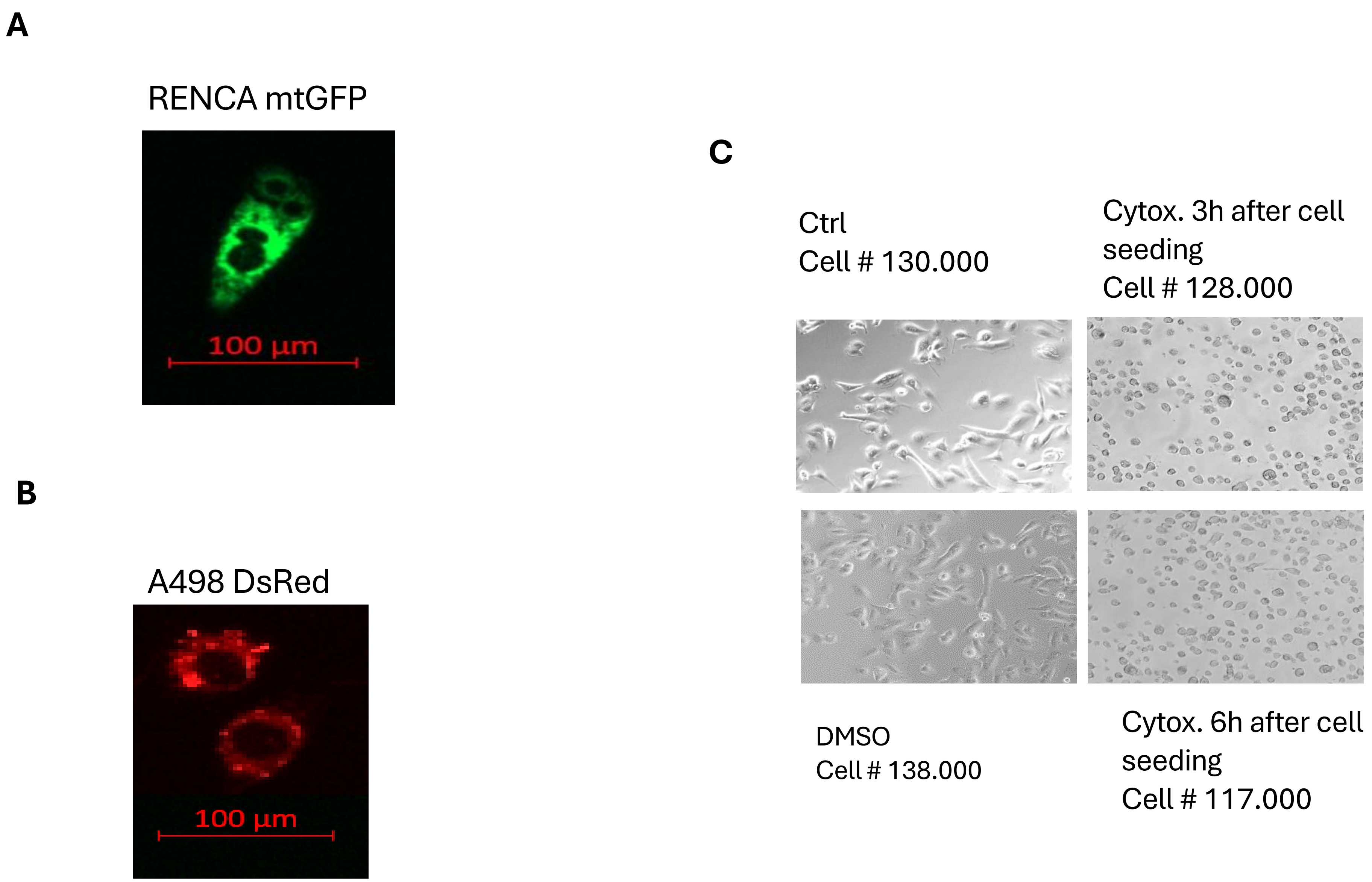


**Supplementary Fig. 3**

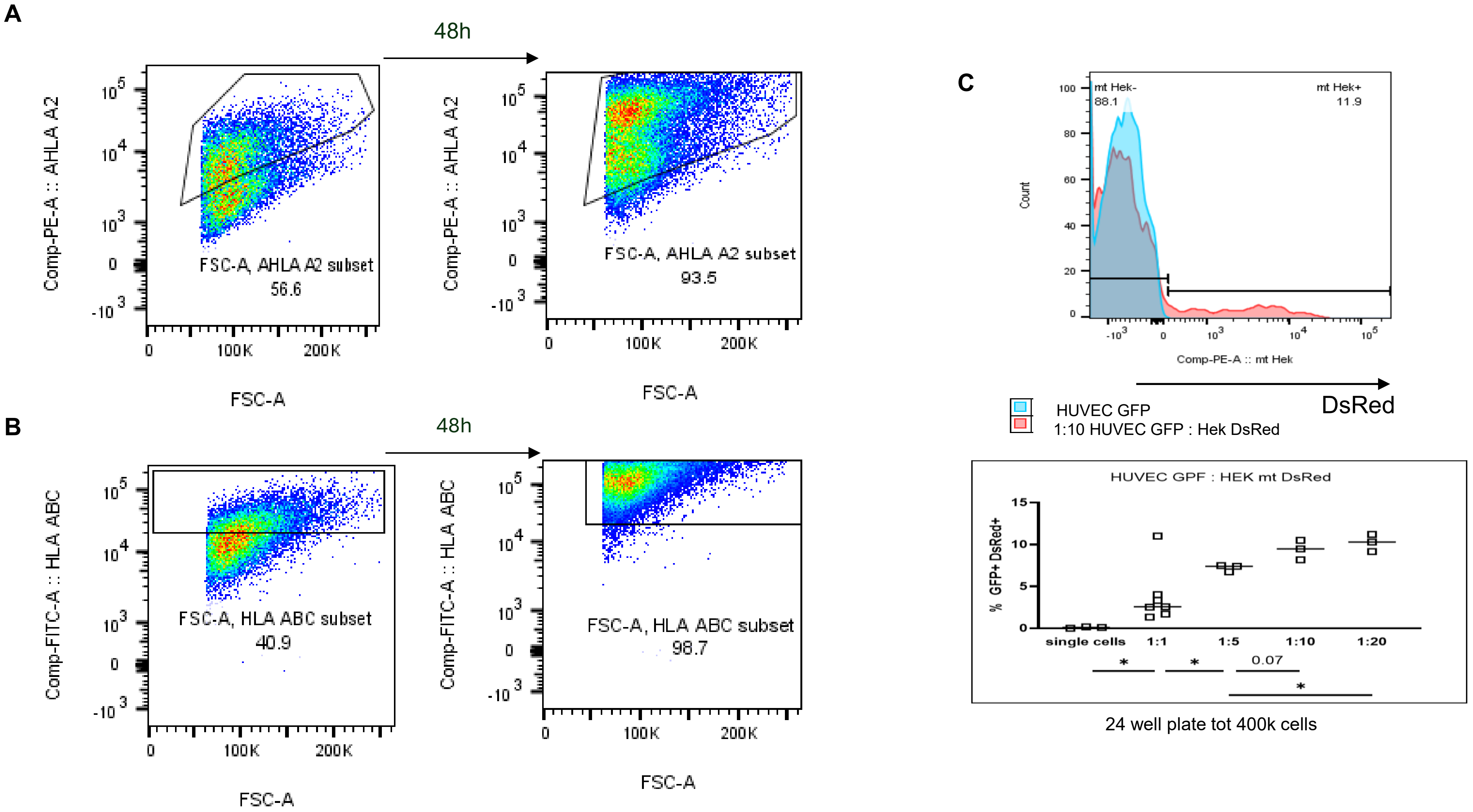


**Supplementary Fig. 4**

| Marker | P value | Hazard ratio | 95% CI |
|---|---|---|---|
| %CD3 | 0.77 | 0.97 | (0.77, 1.21) |
| %FOXP3 | 0.68 | 0.87 | (0.42, 1.81) |
| %CD31 | 0.81 | 1.01 | (0.95, 1.07) |
| COX1 mutation | 0.94 | 1.07 | (0.05, 22.6) |
| CYTB mutation | 0.90 | 0.83 | (0.05, 15.2) |
| ND5 mutation | 0.41 | 0.43 | (0.05, 3.86) |
| mtDNA mutations (0, 1, >1) | 0.44 | NA | NA |
| mtDNA mutations (0, 1–3, >3) | 0.72 | NA | NA |

Multivariable Cox regression (progression-free survival)

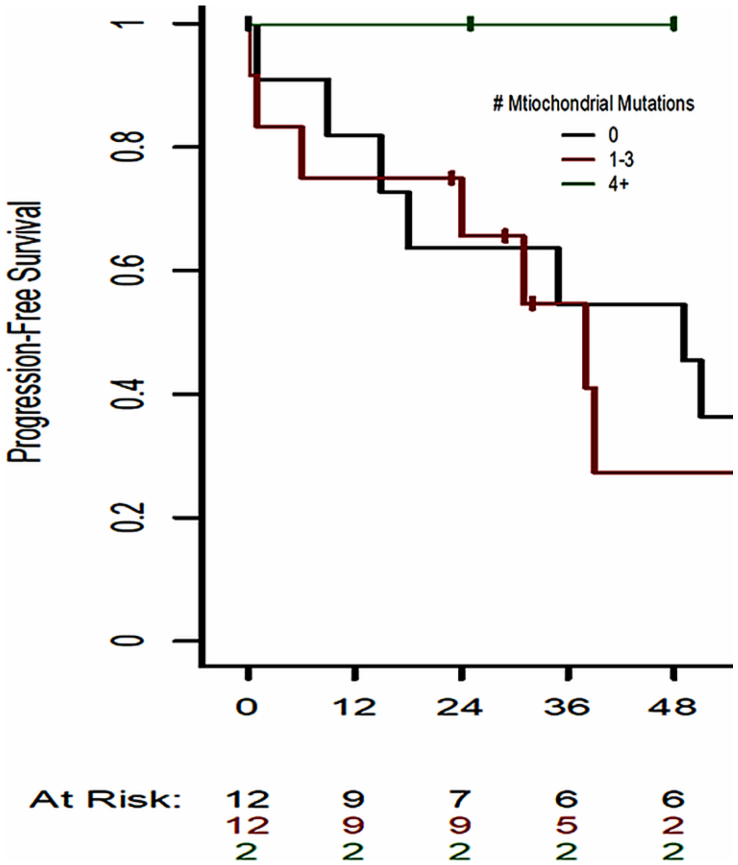


**Supplementary Fig. 5**

### Supplementary Table 1

| Variable | P value |
|---|---|
| Renal vein invasion | 0.59 |
| Grade | **0.002** |
| T stage | 0.16 |
| Histology | 0.79 |

**Supplementary Table 1. Association between mtDNA mutation burden and clinicopathologic variables and outcomes in RCC (n = 30**). Mutation burden showed a significant association with tumor grade (Fisher's exact test).

### Supplemental Table 2

| Endpoint | Spearman ρ | Significance |
|---|---|---|
| %CD3 | −0.045 | n.s. |
| %CD8 | 0.0037 | n.s. |
| %FOXP3 | 0.058 | n.s. |
| %CD31 | 0.135 | n.s. |

**Supplementary Table 2. Association of mtDNA mutation burden metrics with immune and vascular immunostaining endpoints in RCC (n = 52)** Associations between mitochondrial mutational load score and/or mtDNA mutation counts with quantitative immunostaining (%CD3, %CD8, %FOXP3, %CD31) in RCC.

### Supplemental Table 3

| Gene (mutation present vs absent) | %CD3 P value | %CD8 P value | %FOXP3 P value | %CD31 P value |
|---|---|---|---|---|
| ATP6 | 0.6487079 | 0.4869175 | 0.7431576 | 0.31795390 |
| COX1 | 0.6463847 | 0.9141280 | 0.8950380 | **0.04689129** |
| CYTB | 0.8597278 | 0.7438161 | 0.6150679 | 0.49924165 |
| ND1 | N.A. | | | |
| ND2 | N.A. | | | |
| ND4 | 0.7704735 | 0.5917696 | 0.7389469 | 0.94251562 |
| ND5 | 0.5843050 | 0.7616052 | 0.2375422 | 0.062974250 |
| ND6 | N.A. | | | |

**Supplementary Table 3. Association of recurrent mitochondrial gene mutations with immune and vascular immunostaining endpoints in RCC (n = 52).** Recurrent mtDNA gene mutations (present in ≥5 tumors) were scored as present vs absent per tumor and tested for association with immunostaining endpoints.
